# Competing adaptations maintain non-adaptive variation in a wild cricket population

**DOI:** 10.1101/2023.10.14.562337

**Authors:** Jack G. Rayner, Franca Eichenberger, Jessica V. A. Bainbridge, Shangzhe Zhang, Xiao Zhang, Leeban H. Yusuf, Susan Balenger, Oscar E. Gaggiotti, Nathan W. Bailey

**Author notes:** **Corresponding authors:** Jack G. Rayner, Nathan W. Bailey, **Email:**. **Author Contributions:** JGR, OEG, NWB designed and conceived the original study; JGR, FE, JVAB conducted experiments; JGR, JVAB, SZ led data analysis with input from all authors; XZ contributed genomic resources; JGR, LHY, SB, OEG performed fieldwork; JGR & FE performed labwork; JGR led writing of the manuscript with input from all authors. **Competing Interest Statement:** We disclose that we have no competing interests. **Data availability:** Previously unpublished sequencing whole genome, RNA- and RAD-sequencing data are available (or will be once finished processing) in the NCBI SRA under BioProject PRJNA1019311. Scripts used for processing and analysing data are available at https://github.com/jackgrayner/competing_adaptations alongside data collected from mate preference trials, crosses, and life history experiments which will be made permanently available upon acceptance.

## Abstract

How emerging adaptive variants interact is an important factor in the evolution of wild populations. However, the opportunity to empirically study this interaction is rare. We recently documented the emergence of an adaptive phenotype ‘curly-wing’ in Hawaiian populations of field crickets (*Teleogryllus oceanicus*). Curly-wing inhibits males’ ability to sing, protecting them from eavesdropping parasitoid flies (*Ormia ochracea*). Surprisingly, curly-wing co-occurs with similarly protective silent ‘flatwing’ phenotypes in multiple populations, in which neither phenotype has spread to fixation. These two phenotypes are frequently co-expressed, but since either phenotype sufficiently reduces song amplitude to evade the fly, co-expression confers no additional fitness benefit. Numerous negative fitness consequences are known to accompany flatwing, and we find that curly-wing, too, incurs fitness costs via reduced male courtship success and reduced female longevity. We show through crosses, genomic and mRNA sequencing that curly-wing expression is associated with variation on a single autosome. In parallel analyses of flatwing, our results reinforce previous findings of X-linked single-locus inheritance, with the phenotype likely arising through down-regulation of *doublesex*. By combining insights about the genetic architecture of these alternative phenotypes with simulations and field observations, we show that the co-occurrence of these two adaptations impedes either from fixing, despite extreme fitness benefits. Interestingly, both flatwing and curly-wing are statistically associated with nearby inversions, which are also retained as polymorphisms. This co-occurrence of similar adaptive forms in the same populations might be more common than generally considered, and could be an important force inhibiting adaptive evolution in wild populations.

## Introduction

Mutations that confer strong fitness benefits are expected to spread through populations. However, adaptive mutations do not arise in isolation. They interact at the individual level with other fitness-associated alleles in the same genome (1) and at the population level with other segregating alleles. Within a population, multiple beneficial mutations might emerge and segregate contemporaneously. In this case, a given mutation’s adaptive spread will also depend on factors such as its likelihood of recombining into the same genome as other adaptive mutations (2–5). Models of interacting mutations often assume additive fitness benefits in the combined state, suggesting an individual carrying multiple adaptive mutations will have greater fitness than an individual carrying just one. However, this is not necessarily the case, for example if alleles are epistatic (6). Besides epistasis, similar adaptations frequently emerge in lineages evolving under similar selection pressures (7, 8), and can occur through diverse genetic changes even in closely related species (9–12). In such cases it is not obvious that co-expression of alternative adaptations to the same selection pressure should confer any additional fitness advantage.

Here, we investigate this scenario of co-occurring adaptations in a system where multiple phenotypes – adaptive under the same selection pressure, but through a diverse range of morphological changes – have recently emerged across multiple populations of the field cricket *Teleogryllus oceanicus*. Male crickets ordinarily produce song to attract females by rubbing their two forewings together, causing scraper and file structures on opposite wings to make contact (13). However, Hawaiian populations of *T. oceanicus* are attacked by an introduced endoparasitoid fly, *Ormia ochracea*, which uses cricket song to locate hosts for its larvae (14). Researchers have subsequently observed the repeated emergence and spread of multiple song-reducing phenotypes in these populations. First, Zuk et al. (15) observed the emergence of ‘flatwing’ phenotypes, which remove males’ ability to sing via loss or reduction of sound producing structures ordinarily present on the male wing. Flatwing variants have spread through populations on at least three different islands: Kauai, Oahu and Hawaii, but were not observed before 2003. Of the three flatwing phenotypes from across these islands that have been subject to genetic analysis, all are underpinned by X-linked mutations (11, 16–18), but show differing patterns of genomic association that suggest the phenotypes arose independently (11, 16).

We recently documented the emergence of two more reduced-song wing phenotypes, ‘curly-wing’ and ‘small-wing’, in Hawaiian *T. oceanicus* populations (19) (Fig. 1A, Fig. 1B; see also a further protective morph described by (20)). Like flatwing (Fig. 1B), these phenotypes benefit males by reducing or eliminating their ability to sing, which permits them to evade detection by *O. ochracea* (19, 21). A distinguishing feature of curly-wing and small-wing is that both can be visibly expressed by females, whereas the *flatwing* mutation does not affect female wings. Usefully, the three phenotypes are readily distinguishable, allowing us to document their contemporaneous spread. While we have only observed small-wing in two nearby populations, curly-wing – like flatwing – is observed in several populations across the Hawaiian archipelago, perhaps due to recent gene flow (11). Surprisingly, we find that curly-wing and flatwing phenotypes are frequently present in the same populations, and are frequently co-expressed by males. Flatwing phenotypes are present in four of the five study populations in which we observe curly-wing phenotypes, while curly-wing and small-wing co-occur in the fifth (Fig. 1C). The co-occurrence of curly-wing and flatwing is surprising because either phenotype appears to be sufficient to protect males against parasitism (19); flatwing males in particular are almost inaudible in the field. We speculate the co-occurrence of these alternative phenotypes might impede their adaptive fixation, by weakening respective selection coefficients. This would in turn maintain singing-capable phenotypes despite being selectively disfavoured, as the non-adaptive, song-associated variants of either phenotype (i.e., the absence of curly or flatwing morphologies) are shielded from selection when co-expressed with the alternate adaptive variant. Consistent with this, a proportion of audible singing males remain in all populations where two or more reduced-song morphs co-occur, in contrast with two populations in which flatwing alone spread to fixation.

**Figure 1.**
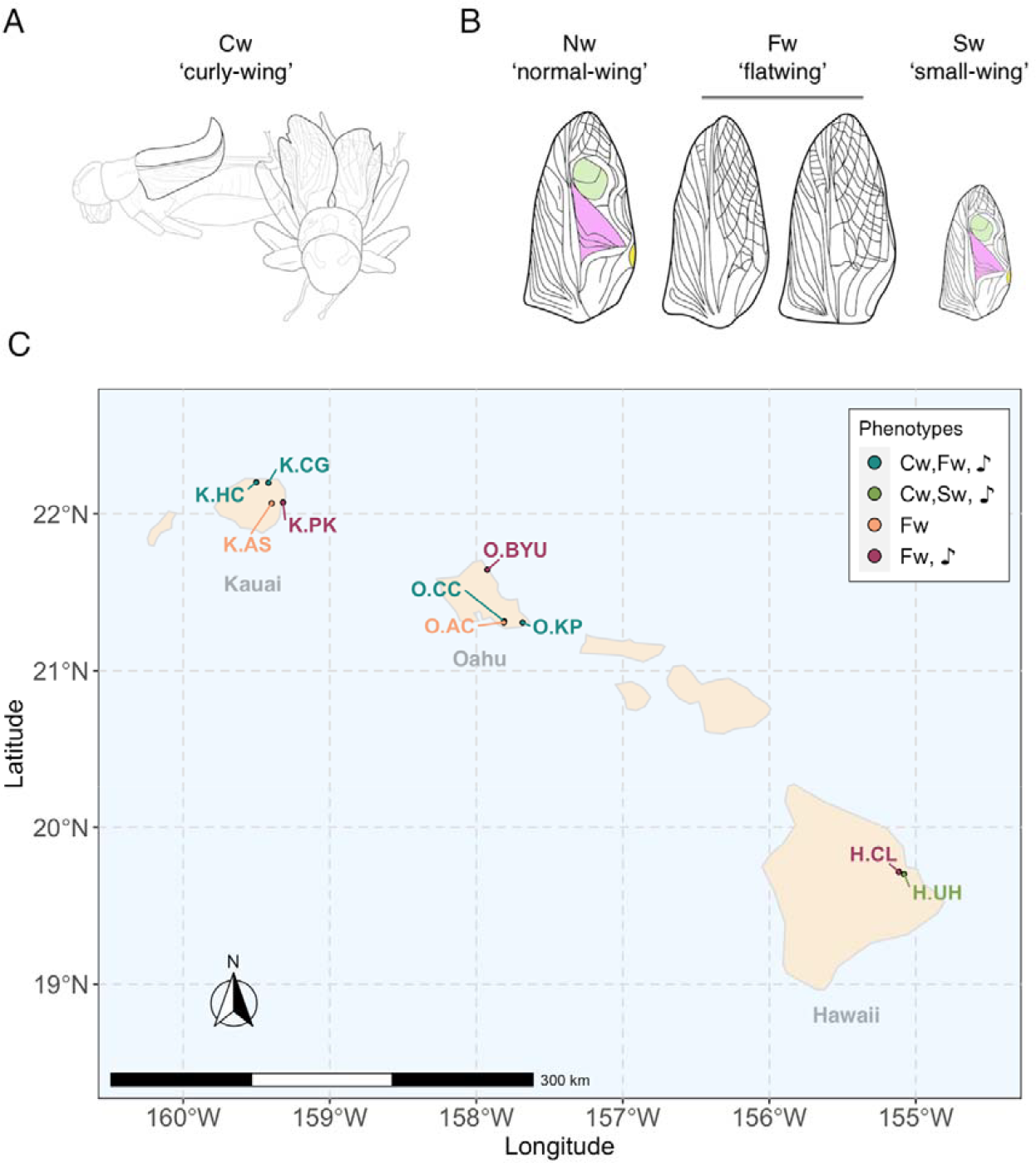
Presence of male-silencing *T. oceanicus* phenotypes across Hawaii. **A)** Side and front-view diagrams of Cw morphology, showing unusually curled forewings, which would ordinarily sit flat. **B)** Diagrams illustrating singing-capable normal-wing (Nw) male wing vein morphology, alongside that of two song-reducing phenotypes: flatwing (Fw) and small-wing (Sw). The two Fw diagrams illustrate variation in Fw morphology between islands and populations in the degree of reduction of sound-producing structures (highlighted in colour). **C)** Distribution of reduced-song phenotypes across the Hawaiian archipelago. In each case, colours indicate which phenotypes are present in each population (i.e., ‘Cw, Fw’ indicates that both Cw and Fw phenotypes are present). ♪ indicates that singing-capable males (i.e., those not expressing any of Cw, Fw or Sw) remain in the population, and this is true of all but K.AgStation and O.AstronomyCenter populations in which Fw has alone spread to fixation.

Here, we evaluate the genetic architecture, fitness consequences, and evolutionary dynamics of co-segregating adaptive variants. The first goal of our study was to assess the heritability of Cw and identify associated genomic regions, as these are as yet unknown. Our second goal was to compare genetic and transcriptomic features of co-occurring curly-wing and flatwing phenotypes, across multiple populations. We present simulations informed by our findings regarding curly-wing and flatwing’s genetic architectures to test our intuition: that co-occurrence of alternative adaptative phenotypes has impeded their adaptive spread in wild *T. oceanicus* populations. Finally, mutant phenotypes which are strongly adaptive in a given context are expected and are often observed to have negative fitness consequences for a range of related and unrelated traits (4, 22). We investigate fitness consequences of curly-wing and flatwing expression in the context of male sexual advertisement, and on adult life history traits of size and longevity in both sexes.

## Results

### Cw is highly heritable and more strongly expressed by females

Half-sibling crosses, performed using laboratory stock originally derived from the Community Center population in Oahu (Oahu.CC), showed curly-wing (Cw) is highly heritable and segregates in a manner consistent with autosomal inheritance (Fig. 2A). In a linear mixed model, a random effect term including parental phenotypes and identities explained an estimated 90% of variance in the proportion of Cw offspring (est. R^2^ of random effects=0.903). Replacing our response variable with average offspring curliness score (Fig. S1, Table S1) slightly reduced the estimated proportion of variance explained (est. R^2^ of random effects = 0.878), so we treat Cw as a discrete trait. Fifty-eight percent of males used in crosses expressed flatwing (Fw), and patterns of inheritance were consistent with X-linkage previously observed for Fw variants (16, 17) (Fig. S2). Among male offspring, there was no correlation between expression of Cw and Fw phenotypes, supporting the view that causal regions are located on different chromosomes (Table S2), which is likely to be a key feature of their evolutionary interaction.

**Figure 2.**
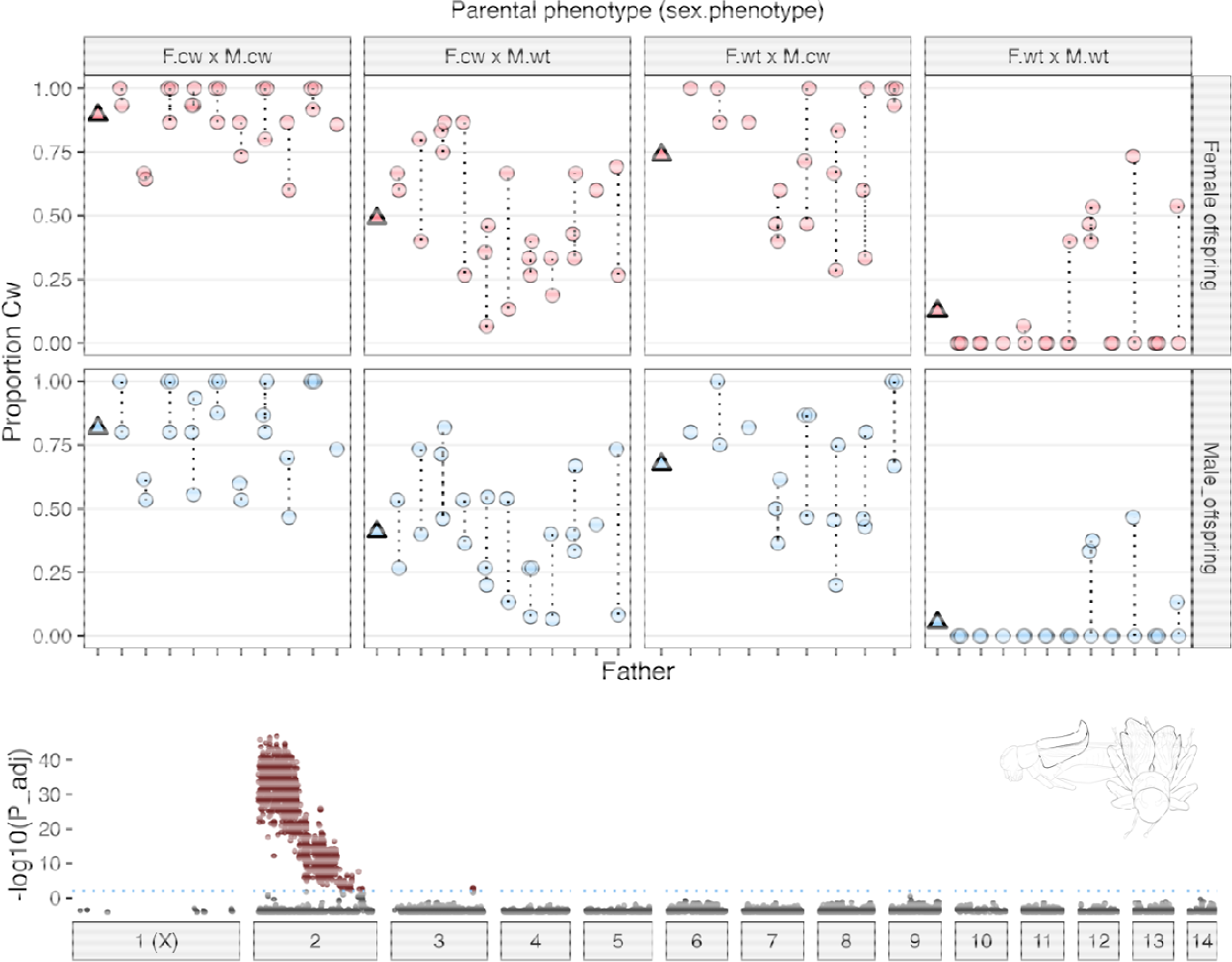
Inheritance and genetic architecture of Cw. (**A**) Each circle shows the proportion of full-sib F_1_ offspring expressing curly-wing following mating of the respective F_0_ male (X-axis) with a single female. Dotted lines connect half-sibling families. Panels are separated into groups based on parental curly-wing phenotype (F.cw x M.cw = female curly-wing mated with male curly-wing, and so on), and by offspring sex. Triangles show mean values for each panel. **(B)** Genome-wide association significance (-log_10_ Bonferroni-adjusted P) between allelic variants and Cw phenotype.

Although an overwhelming proportion of variance in curliness observed between families was explained by parental ID and phenotype, curliness was also associated with offspring sex (X^2^_1_ = 24.816, P<0.001) and rearing density (X^2^_1_ = 8.107, P = 0.004; full model est. R^2^ of 0.932). Females and offspring reared at lower density were each more likely to express Cw, and females were assigned higher curliness scores (females: x□ = 3.86 ± 0.003 SE, males: x□ = 3.06 ± 0.003 SE). Sex and rearing density also influence expression, in the same direction, of an analogous curled wing phenotype, ‘curly’, in *Drosophila* (23, 24). Across field cricket taxa, sex and rearing density each affect development rate (25, 26), suggesting a plausible explanation is that wing curliness is influenced by growth rate, as has been found in *Drosophila* (27).

As expected in the case of Cw being underpinned by one or a few co-localised major effect loci, Cw × Cw and Wt × Wt crosses produced predominantly Cw and Wt offspring, respectively (Fig. 2A), albeit with exceptions attributable to incomplete penetrance. In crosses where just one parent visibly expressed Cw, paternal phenotype was more strongly correlated with the proportion of Cw offspring. The cause of this parent-of-origin effect is not clear, as inheritance patterns do not support sex-linkage, but it is plausible that Cw penetrance is influenced by one or more X-linked alleles, which might also contribute to sex differences in expression.

### Cw is genetically associated with a single autosome, Chromosome 2

We performed genome-wide association tests between 178 Cw vs. 197 Wt individuals from a single inbred F_2_ family, with 13,822 filtered RAD markers mapped to the *T. oceanicus* reference genome V2 (Zhang et al. *in review*). This family was produced from laboratory stock derived from the wild Oahu.CC population. Phenotype-genotype association tests detected significantly Cw-associated variants across nearly the full length of the 245 Mb Chromosome 2 (Chr2), with the association strongest across the first ca. 85 Mb (Fig. 2B). While there were no discrete peaks within this region, inspection of heterozygosity and genotypes across Chr2 highlighted a region between 66:83 Mb showing consistent differences between Wt and Cw phenotypes (Fig. S3).

Chr2 also showed elevated numbers of differentially expressed (DE) genes associated with Cw (Fig. S4). Fifty-two genes were DE between developing wings of Cw vs Wt males (DE_Cw_), whereas 225 were DE between those of Nw and Fw males (DE_Fw_). In both cases, DE genes were concentrated on chromosomes harbouring the respective causative variants, particularly for DE_Cw_ genes, 50% of which were located on Chr2 (Fig. S4). This represents strong overrepresentation given that Chr2 accounted for just 12.2% of the 12,049 genes present in the filtered transcriptome (X^2^_1_ = 64.707, P < 0.001). Moreover, 24 of these 26 were within the 0:85 Mb region strongly associated with Cw in the RAD-seq data.

To perform tests of functional enrichment of gene ontology (GO) terms among differentially expressed genes, we first relaxed the threshold for the DE_Cw_ gene set to FDR < 0.1 to increase our statistical power. Of 98 such genes, a similarly high proportion (N=45; 46%) were located on Chr2, indicating this subset was biologically meaningful. Of 52 genes that could be assigned GO terms, one molecular function (*endopeptidase inhibitor activity* [N = 4 genes, versus 24 in the transcriptome; binomial test P_adj_=0.033]) showed evidence of enrichment. This was driven by serpin genes, homologous with *Spn55B, SPn42Dd* and *Spn43Aa* in *Drosophila*, which were enriched 22-fold. In insects, serpins play a primary role in innate immunity via the melanisation cascade (28), but have also been repeatedly implicated in abnormal wing development. In *Drosophila, Serpin88Ea* is expressed in developing wings, and knockdown results in defective unfolding and wing expansion following the adult moult, with the adult phenotype showing similarity with curly-wing (29). A similarly altered wing phenotype, again showing visible similarity to curly-wing, was observed following knockdown in *Drosophila* of another serpin gene, *Serpin-27A* (30).

Among the 225 DE_Fw_ genes, 50 (22.25%) were located on the X chromosome (cf. 12.0% of all genes in the filtered transcriptome; X^2^_1_ = 20.616, P<0.001) (Fig. S4). Two were 14Kb and 3Kb upstream of the annotated *dsx* gene (259.05 – 259.44 Mb), the latter of which showed significant blastx homology with ‘doublesex and mab-3 related transcription factor’ in the desert locust (*Schistocerca gregaria*). Both were down-regulated in Fw males, consistent with down-regulation of *dsx* in developing Fw wing tissue observed by Zhang et al. (11), but at a later stage of tissue development and in a population not previously examined.

### Evidence for a large Cw-associated inversion

The large Cw-associated region between on Chr2, without an obvious peak of genetic association (Fig. 2B), could implicate a large structural variant such as a chromosomal inversion. Using existing whole genome sequencing (WGS) data collected in 2017 from the wild Oahu.CC population, the same individuals from which our lab stock was derived, we found strong evidence of a segregating inversion in this region. Specifically, principal component analysis (PCA) separated samples into three discrete clusters: representing homozygotes for the two divergent non-recombining haplotypes, and an intermediate cluster of heterozygote samples (31) (Figs. S5, S6). Samples from these clusters showed drastically different rates of heterozygosity and strong patterns of linkage within the first 80 Mb of Chr2, consistent with expectations under the scenario of a segregating inversion (32, 33) (Figs. S5, S6). We used *svdetect* to predict breakpoints associated with large (> 5Mb) inversions in this region based on paired-end read alignments (34). After filtering for location and expected frequencies across samples based on PCA clustering, there remained a large predicted inversion corresponding to our observations between 7.5 and 80 Mb on Chr2. This approach has a high false positive rate with WGS data, but for convenience we henceforth treat the inversion as located within the region of 7.5—80 Mb on Chr2.

### Phenotype-genotype associations of similar morphologies across populations

Cw and Fw phenotypes show surprising geographical overlap in their occurrence across *T. oceanicus* populations. To investigate the genomic architecture of Cw and Fw phenotypes in share populations, and extend our analysis of Cw across multiple populations, we examined WGS data collected in 2021/2022 from males of known phenotype from the focal Oahu.CC population (13 Cw, 17 Wt; 8 Fw, 22 Nw), alongside populations from two other Hawaiian islands: Kauai.CG (19 Cw, 11 Wt; 16 Fw, 14 Nw), and Hawaii.UH (18 Wt, 12 Cw, all Nw) (see Fig. 1). Neither Cw nor Fw phenotypes have reached fixation in any of these populations, and in Oahu.CC and Kauai.CG, 7 and 12 males, respectively, co-expressed Cw and Fw. Given prior knowledge regarding their genetic architectures, we focussed analyses of Cw and Fw phenotypes on Chr2 and the X, respectively. For analyses of Fw, the Oahu.CC data was combined with data collected from 2017 for a total sample of 50 (18 Fw, 32 Nw).

We found a strong association between Cw and genetic variation across the region of the predicted inversion at 7.5:80 Mb on Chr2. The pattern of clustering across populations was again strongly consistent with the presence of an inversion in this region, separating samples from all populations into three discrete clusters on PC1 (Fig. 4A). Moreover, the inferred frequency of the inverted haplotype differed significantly between Cw and Wt samples in all three populations (Wilcoxon rank-sum test: P_Hawaii.UH_=0.005; P_Kauai.CG_=0.005; P_Oahu.CC_=0.011), however, the pattern of Cw – PC1 association in Hawaii.UH was opposite that of Kauai.CG and Oahu.CC. To investigate this incongruity, we performed an F_ST_ scan (10 kb window, 10 kb step size, using Weir and Cockham’s F_ST_ implemented in vcftools (35)) between samples inferred to be homozygous for the inverted haplotype in Kauai.CG and Hawaii.UH, but which expressed opposite phenotypes (Fig. 4A). F_ST_ values of up to 1 were identified in three regions of striking genetic divergence: ca. 7.5:15 Mb, 60:70 Mb, and 80 Mb (Fig. 4B). Visualisation of linkage and heterozygosity along Chr2 also revealed the Hawaii.UH population showed quite different patterns in the region of 7.5:80 Mb compared with Oahu.CC and Kauai.CG populations (Fig. S7). This might suggest recurrent chromosomal rearrangements in this region, a pattern that seems to be relatively common (33, 36, 37). We anticipated that the Cw-associated variant(s) might sit in one of these three windows showing high F_ST_, as variants in these windows are likely to be statistically associated with the inversion in each population via linkage, but could show opposite patterns of association between populations (i.e., gametic coupling in Kauai.CC and Oahu.CC, but repulsion in Hawaii.UH).

**Figure 3.**
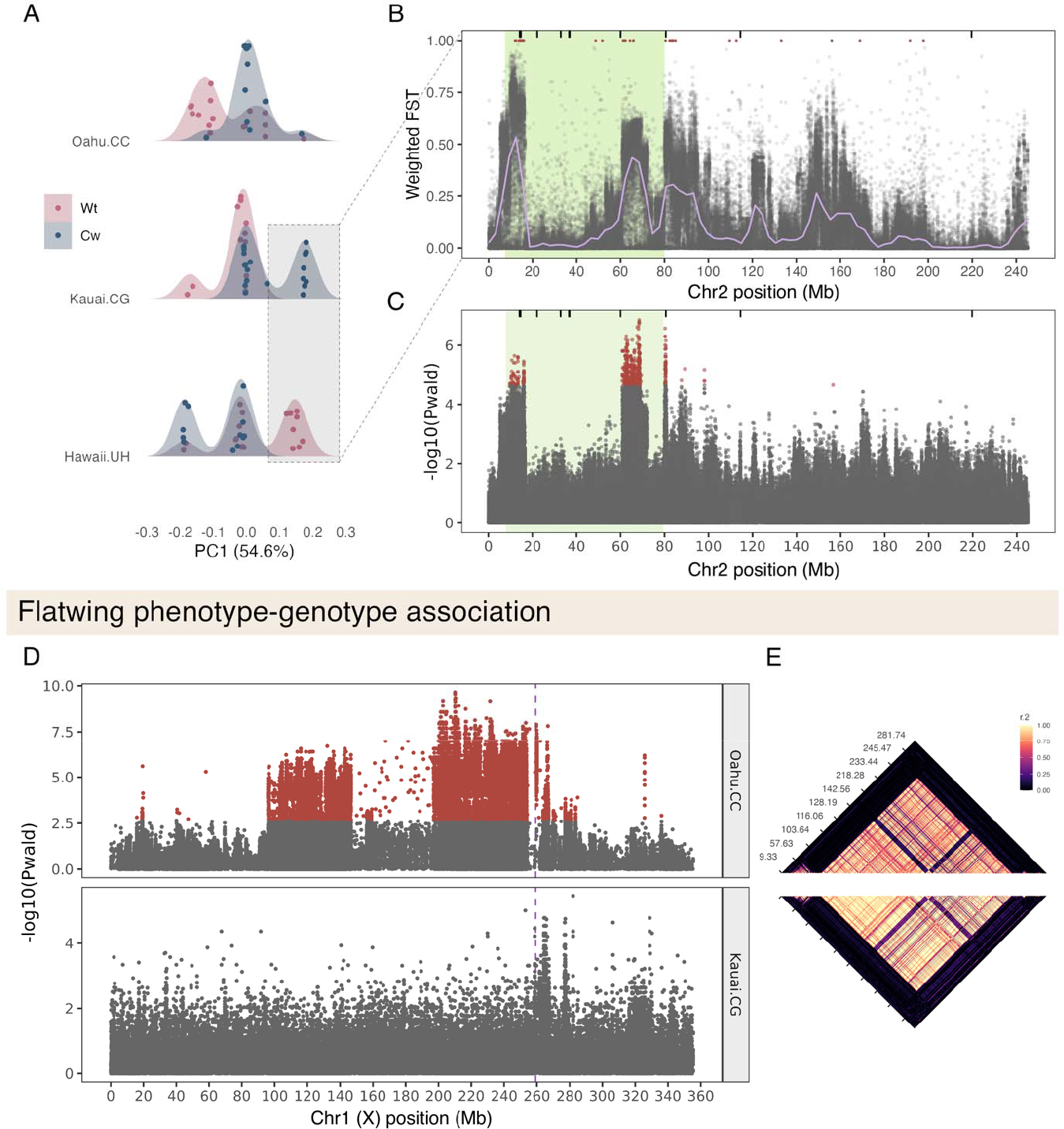
Genomic regions associated with Cw and Fw. **A)** PC1 clustering across populations from PCA analysis of SNPs within 7.5 Mb and 80 Mb regions of Chr2. Points are jittered on the Y-axis to fill the density distribution illustrated by curves, and both are coloured by sample phenotype. **B)** Weighted F_ST_ values between Kauai.CG and Hawaii.UH populations along Chr2, using samples in the highlighted rectangle from (A). Windows with an F_ST_ of 1 are highlighted in red. **C)** SNP-wise association tests between Cw and Wt phenotypes across all populations, with a single SNP with Bonferroni-adjusted P < 0.05 in red. Vertical lines on the top of the plots B and C show the locations of genes with an annotated function in serine-type endopeptidase inhibitor activity, and the region of the putative inversion is highlighted in green. **D)** SNP-wise association tests between Fw and Nw phenotypes within Kauai.CG and Oahu.CC populations, on the X chromosome. The dashed purple line indicates the position of *dsx*. **E)** Linkage across the full X-chromosome in Oahu.CC (top) and Kauai.CG (bottom).

**Figure 4.**
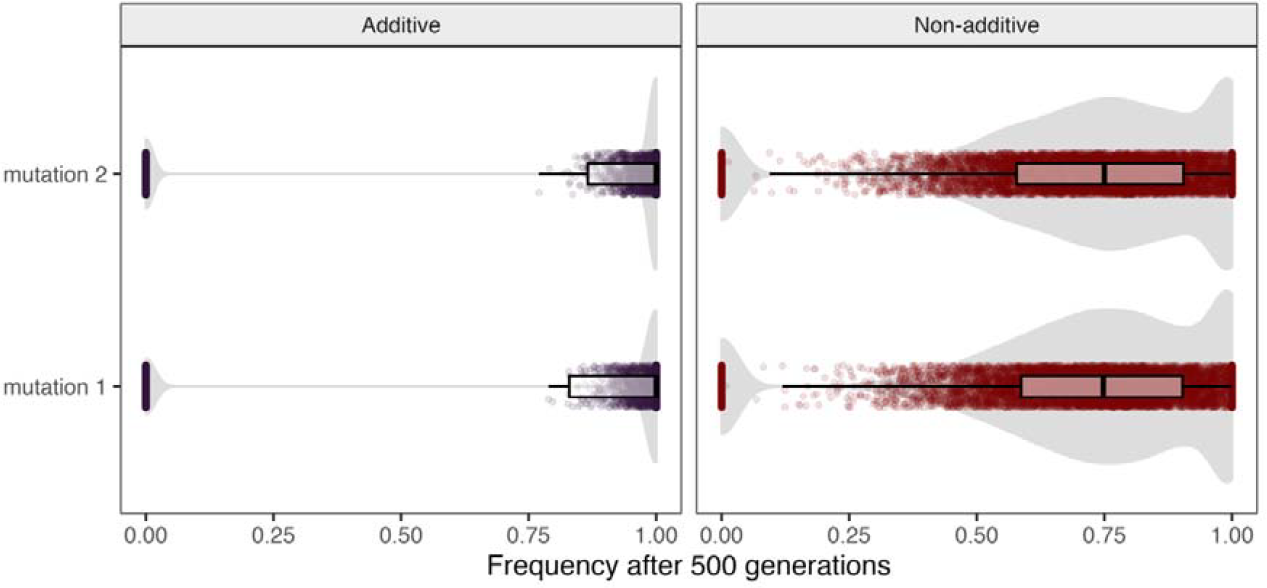
Non-additive fitness benefits of competing mutations impedes their spread. Frequencies of two mutations with equal fitness benefits after 500 generations of evolution under simulated scenarios of additive and non-additive fitness benefits in the co-expressed state. Boxplots show medians and interquartile ranges. Points are randomly jittered on the Y-axis for interpretability, and the grey filled areas illustrate their density distribution.

In an association test between Cw and Chr2 SNPs across samples from all three populations (N = 1,438,882 SNPs), performed using GEMMA (38), several SNPs reported a significant (P_adj_ < 0.05) association with Cw (Fig. 4C). All were within or immediately adjacent to the putative inversion in Kauai.CG and Oahu.CC, and nearly all were within candidate regions highlighted by the F_ST_ analysis above. We observed strong linkage between the 100 top SNPs (Fig. S8, S9), impeding our ability to identify candidate regions. While these SNPs showed clear divergence between Cw and Wt phenotypes (Fig. S9), there were also exceptions; in particular, four of the 46 Wt samples were consistently genotyped for Cw-associated variants, which we expect is due to the incomplete penetrance of the Cw mutation(s) previously observed (Fig. 2A). Five of these SNPs were located within annotated genes. These included a SNP at 80,625,117 bp within *ITIH4*, which has an annotated function in endopeptidase inhibitor activity (GO:0004867; highlighted by the RNA-seq analysis and by involvement in Cw-like mutant phenotypes in *Drosophila*), and sits within a region containing several DE_Cw_ genes (Fig. S10).

Flatwing phenotypes have been repeatedly associated with *dsx* at genomic and transcriptomic levels (11). We investigated Fw-associated variants in Oahu.CC and Kauai.CG (N = 997,680 and 587,058 X-linked SNPs, respectively). We analysed each population separately as prior analysis have suggested Fw variants from these islands have independent genetic architectures. In Kauai.CG, none of the most strongly associated SNPs reached statistical significance at P_adj_ < 0.05, probably in part due to the small sample, whereas in Oahu.CC, there were strongly Fw-associated SNPs across nearly the full range of X chromosome (Fig. 4D). In both populations, we observed strong linkage between SNPs across a very large portion of the X chromosome (Fig. 4E). PCA of X-linked variants led samples to group into two non-overlapping clusters, shared between populations, supporting the presence of another, extremely large, inversion. This may have contributed to the low density of RAD SNPs on the X in our inbred mapping family (Fig. 2B). Fw and Nw samples were not separated on PC1 or PC2 in either population (Fig. S11), indicating the inversion is not causally associated with Fw – though there was a statistical association between the inversion and Fw in Oahu.CC (Wilcoxon rank-sum test: P = 0.001). Inspection of linkage patterns in both populations suggested the inversion spans the region of ca. 95.7 – 253.2 Mb, so we considered the top 50 Fw-associated SNPs outside of this region from each populations None of these were shared between populations. However, 39 (78%) of these top 50 SNPs in the Oahu.CC population were within 1 Mb of *dsx* (259.05 – 259.44 Mb), whereas in Kauai.CG one SNP, at 258,733,070, was ca. 320 Kb from *dsx*. These SNPs showed consistent and strong, but imperfect genotypic divergence between Nw and Fw samples in each population (Fig. S12). This potentially indicates causative Fw mutations are near to but not included in this set of SNPs, which could be the case if Fw is caused by an indel, or variant otherwise filtered from our dataset. Generally speaking, our results support those of (11, 16) in finding no overlap in the most strongly Fw-associated SNPs between populations, but repeatedly highlighting the region surrounding *dsx*.

### Interacting mutations under conditions of additive benefits and redundancy

Informed by the non-overlapping genetic architectures of the two phenotypes, we evaluated the prediction that Cw and Fw segregating in the same population would impede either from reaching fixation, by running simulations in SLiM v4.0.1 (39). The results supported our prediction. In a single population of 500 diploid individuals, we introduced two dominant mutations *m1* and *m2*, each, on unlinked chromosomes in a separate random subset of 5 genomes. Simulations were run under two conditions: in the ‘additive’ scenario, the mutations each conferred an additive fitness benefit of 0.15, meaning a combined fitness benefit of 0.3 when co-expressed. In the ‘non-additive scenario, the mutations each conferred a fitness benefit of 0.3, unless co-expressed, in which case they each conferred a fitness benefit of 0.15 (resulting in a combined fitness benefit of 0.3). We chose these parameters because they result in equal fitness benefit across scenarios when the two mutations are co-expressed. Supporting our intuition, we found across 10,000 simulations that both mutations were much more likely to be segregating at intermediate frequencies after 500 generations in the non-additive scenario. In the additive scenario, both mutations were still segregating after 500 generations in just 6.44% of simulations, versus 76.93% in the non-additive scenario (Fig. 5): in which, as for Cw and Fw, the two mutations do not confer additive fitness benefits.

**Figure 5.**
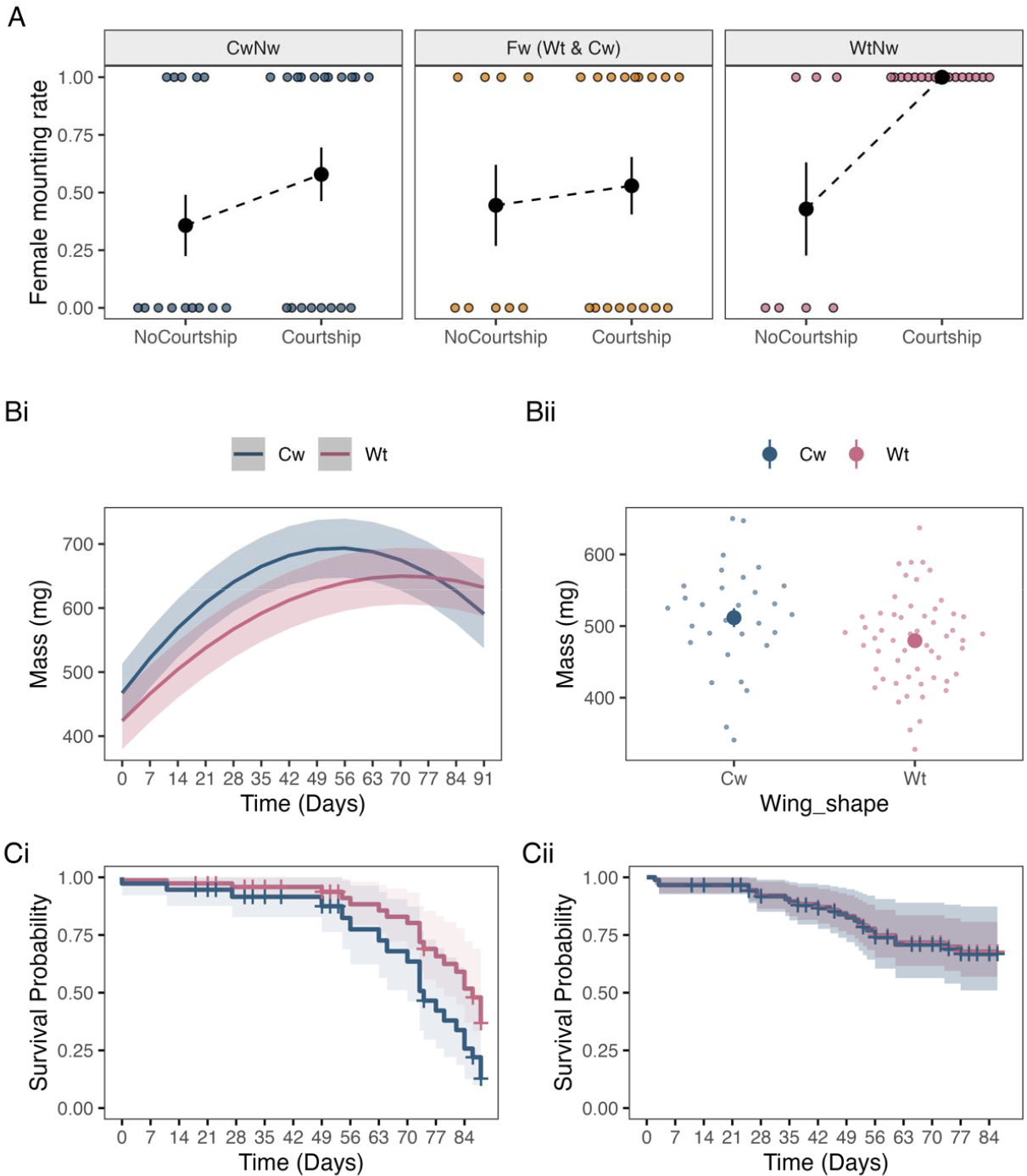
Correlated fitness consequences of adaptive mutations. **A)** Effects of courtship (measured by attempt to produce courtship song) by males of different wing phenotypes upon rates of female mounting (i.e, decision to mate). Black points and error bars show means ± SE. **B) (i)** Predicted mass ± standard error across adult ages in females. **(ii)** Observed mass at 14-days post-eclosion for males, with solid points showing means ± standard error. **(iii, iv)** Probability of survival (solid lines) at adulthood based on proportional hazards regression of Wt and Cw phenotypes in females and males, with shading indicating 95% confidence intervals.

### Correlated fitness consequences

Although Cw and Fw do not appear to combine to confer greater fitness benefits in the co-expressed state, mutations that spread under strong selection frequently also incur fitness costs through direct or indirect pleiotropic effects and genetic hitchhiking. Such negative fitness consequences are more likely in cases where large effect mutations are favoured due to extreme displacement of a population from a fitness optimum (4, 40), as is the case in *T. oceanicus* populations parasitized by *O. ochracea* (18, 41), and would likely combine additively. As a result, males expressing both phenotypes could actually be disadvantaged, and female carriers might also suffer negative fitness-associated consequences of Cw and Fw, despite male-limited fitness benefits (females being obligately silent).

In grylline field cricket species such as *T. oceanicus*, females exert control over mating interactions as they must mount males in order for mating to occur. Male song is an important courtship trait, and males that can sing have a strong advantage in mating interactions (21). In mating trials, we found female mate choice differed predictably between song-producing normal-wing (WtNw), flatwing (WtFw and CwFw) and Cw (CwNw) phenotypes in accordance with their varying ability to produce courtship song (binomial GLM of female decision to mount: MaleMorph × Courtship X^2^_2_ = 7.54, P = 0.023). As expected, males expressing Cw or Fw morphology were strongly disadvantaged in the context of courtship, compared with WtNw males for which courtship consistently elicited female mounting (Fig. 6A). Including repeated measures from replicate trials that re-used previously-tested individuals produced qualitatively similar patterns, indicating their reliability (Fig. S12), but impeded model fit after inclusion of random intercepts for male and female ID. While the fitness cost associated with reduced courtship ability is evidently substantial, though the spread of songless phenotypes in *T. oceanicus* across the Hawaiian archipelago, in some cases to fixation (19, 42), suggests this cost is secondary to fitness benefits gained from evading detection by *O. ochracea* (21).

Apart from direct effects of reduced signalling ability on male fitness, adaptive mutations might also exert pleiotropic, or other indirect fitness consequences. Most strikingly, females expressing Cw morphology had significantly reduced longevity (P<0.001; Table S3), providing evidence of strong negative fitness effects of the phenotype in females, who also do not benefit directly from loss of song. Neither Cw nor Fw phenotypes appeared to affect male longevity, though we recorded low male mortality in general over the course of the experiment (27.3% dying in the 91 days post-adulthood, vs 45.7% in females) (Table S3; Fig. 5). Similarly, male mass at 14 days adulthood was unaffected by wing shape (Cw vs. Wt) or wing venation (Nw vs. Fw) (Table S4). In females, however, Cw interacted non-linearly with adult age in affecting mass; Cw females typically had greater mass, but this effect diminished at later ages. This pattern was also largely reflected in scaled mass index, sometimes used as a measure of body condition (43). Fitness consequences of greater body mass in female carriers of Cw are thus unclear, contrasting with clear fitness costs demonstrated by reduced longevity. (Fig. 5; Table S3-S5)

## Discussion

Evolutionary dynamics following the contemporaneous emergence of multiple beneficial mutations have long attracted interest, but opportunities to observe such dynamics empirically are rare – exceedingly so in wild populations. This limitation does not exist for the Hawaiian field cricket populations we studied. Under extreme selection against song, two different adaptive song-loss phenotypes, Fw and Cw, have repeatedly spread in *T. oceanicus*. We find these two phenotypes have non-overlapping genetic architectures, and are frequently co-expressed despite their lack of additive fitness benefits when combined. Our findings suggest the co-occurrence of these two phenotypes, either of which is sufficient to protect males from parasitism, has impeded the fixation of either causative mutation.

Adaptive mutations are expected to be very rare (44). This has clearly not been the case among typically small and fragmented Hawaiian populations of *T. oceanicus* evolving under fatal parasitism by *O. ochracea*, across which four different and apparently novel song loss ‘morphs’ have emerged within the last 20 years (15, 19, 20). On top of these divergent wing morphs, superficially similar flatwing phenotypes have apparently arisen independently on at least three occasions (11, 16). The reason for this exceptional proliferation of adaptive forms seems clear. The introduction of *O. ochracea* to Hawaiian populations of *T. oceanicus* radically changed the fitness landscape, from one in which male singing ability is strongly favoured by benefits in attracting female mates, to one in which any mutation that corrupts a male’s ability to sing offered a selective advantage. While this scenario of similar adaptive mutations competing within a population might be presumed to be rare, we point to the widely observed potential for similarly adaptive phenotypes to emerge and spread across populations and species (i.e., parallel, convergent, or repeated evolution (45)), at least some of the time through different genetic mutations (8). The opportunity for alternative adaptive mutations to be introduced to the same populations through either, or a combination, of gene flow and mutational input might therefore be underappreciated, and frequently overlooked.

The maintenance of non-adaptive variation underlying singing ability in male crickets is of particular evolutionary importance because silent male *T. oceanicus* benefit from the retention of singing males, which they rely upon to adopt satellite mating tactics (15). Additionally, cricket song plays an important role in the social environment, affecting a range of traits such as adult reproductive investment (46), neural gene expression (41), and locomotive activity (47). Should selection against song decrease in severity (e.g., via *O. ochracea* population decline) in populations in which co-occurrence of Cw and Fw phenotypes has impeded the loss of variation underlying singing-capable male phenotypes, the latter would be expected to once more spread through the population. Males would thus regain the ability to attract female mates, without requiring the secondary evolution of song or other signalling modalities (48, 49).

While Cw and Fw do not combine additively in the context of fitness benefits, non-adaptive consequences of the two phenotypes could combine additively to reduce net fitness benefits, further impeding their spread. We found evidence of correlated fitness consequences of the curly-wing phenotype, particularly in females, who suffered greater adult mortality and exhibited greater mass, while males suffered reduced courtship ability. Size and longevity, assayed here, are only a subset of many traits that might be impacted by negative pleiotropy or genetic hitchhiking associated with a strongly adaptive mutation. While we did not observe an effect of flatwing upon male longevity or size in the Oahu.CC stock population, previous work has found flatwing variants are associated with various phenotypic effects including altered male reproductive investment (25, 46, 50, 51); accelerated growth rate (25); and reduced female fecundity (52). Differences in the magnitude of fitness costs associated with either phenotype would also lead to different selection coefficients acting upon each phenotype. However, this would be challenging to quantify systematically, and in populations such as those of Hawaiian *T. oceanicus*, which are typically small and fragmented, might have little influence upon long-term evolutionary dynamics.

Standing genetic variation is of central importance to understanding the ability of wild populations to adapt under extreme selection, particularly when selection is strong and adaptation must occur quickly (53). However, selective sweeps erode genetic variation. In wild populations of *T. oceanicus*, we find phenotypic and genetic variation in the form of multiple, contrasting wing morphologies has been maintained despite extreme selection for reduced song amplitude. In addition, flatwing and curly-wing phenotypes are each statistically associated with large, polymorphic inversions. This maintenance of variation in the face of strong selection is due, at least in part, to non-additive fitness benefits of interacting adaptation; however, rather than facilitate adaptive evolution by maintaining variation, competition between alternative adaptations impedes their fixation. Our findings demonstrate that the interaction between selection and genetic variation can be difficult to predict, highlight distinctive evolutionary dynamics when similar adaptations co-occur, and illustrate the importance of studying adaptive evolution prior to the fixation of adaptive variants.

## Materials and Methods

Full details of methods are provided in the supporting information.

## Acknowledgments

We thank landowners in Hawaii for permission to collect crickets on their grounds, and gratefully acknowledge the Natural Environment Research Council (NE/T0006191/1) for funding to NWB, OEG & JGR that supported this study. We acknowledge computational resources and generous technical support from the James Hutton Institute Bioinformatics HPC (BBSRC grant BB/S019669/1). We thank Tanya Sneddon, Audrey Grant, Megan McGunnigle, and David Forbes for assistance with laboratory work, Ana Drago Rosa and Renjie Zhang for assistance with fieldwork, and Michael Ritchie, Renjie Zhang, and Thomas Hitchcock for useful comments, discussion, and feedback in preparing the manuscript.

## Notes

### Competing Interest Statement

The authors have declared no competing interest.

