## Supporting Information for "Competing adaptations maintain non-adaptive variation in a wild cricket population"

### Supporting methods

#### Forewing nomenclature

All wing phenotypes discussed are expressed in forewings, which are used by crickets to sing, whereas hindwings are involved in flight. At the level of 3-dimensional wing morphology, we refer to curly-wing (Cw) and wildtype (Wt) phenotypes, where Wt indicates the absence of Cw morphology (cf. Fig. 1A). By contrast with Cw, Wt forewings lie flush, right overlapping left, on the dorsal surface of the abdomen when at rest. We also refer to normal-wing (Nw) and flatwing (Fw) wing vein morphology, using nomenclature established by Zuk et al. (2006). Flatwing and curly-wing are therefore not opposite, but instead describe different male-silencing phenotypic morphs which can be co-expressed (CwFw). Using this nomenclature, typical singing males are WtNw, whereas any of WtFw, CwNw or CwFw combinations exhibit strongly reduced or absent singing ability (Table S7). Note that Fw is sex-limited, in that it cannot be detected by eye when females carry causal genetic variants. In contrast, Cw can be expressed by both males and females.

#### Heritability crosses

We performed half-sibling crosses between 37 males and 111 females, for each of the four possible combinations of parental wing phenotype (Cw/Wt) ⨉ parental sex (M/F). Each male was sequentially paired with three females, for five days each, to produce up to three half-sib F_1_ families. In each of these combinations a mix of normal-wing and flatwing male phenotypes were included, with female genotype at the *Flatwing* locus unknown. Offspring number was recorded 30 days after hatching for each pairing that produced offspring, and where necessary numbers were then reduced by culling to avoid overcrowding (<~150 juvenile crickets per 20L box). The curliness of each parental male and female’s wings was scored according to a qualitative scale (0-4 per wing; Fig. S2; Table S1).

Offspring from each of the successful crosses were reared to adulthood and the curliness of each of their wings was similarly scored on discrete (Cw vs. Wt) and qualitative scales. We aimed to record the phenotype of >=15 of each sex from each full-sib F_1_ family. From one of these F_1_ families, five male/female pairings with contrasting curliness scores (one Wt, one highly Cw) were removed and paired to produce inbred F_2_ families, one of which was used for association mapping, and another which was used for RNA-seq analysis.

Results of crosses were analysed using generalised mixed models with a response variable of mean offspring Cw phenotype. We included random intercepts for maternal ID nested within paternal ID, nested within group (the latter referring to the four combinations of parental sex ⨉ phenotype [Cw/Wt]), i.e. (1|grp/father/mother). Models included fixed effects of rearing density in the first 30 days of development and offspring sex. We ran the model with both quantitative (mean offspring Cw scores) and qualitative (proportion of Cw offspring) responses, using gaussian and binomial error distributions, respectively. Models were fit using lme4, tested using type II Wald’s Chi-square tests in R, and R-squared values were estimated using the R package MuMIn (54).

#### Genetic mapping

We performed restriction site associated DNA sequencing (RAD-seq) to obtain sequences of single-end DNA distributed across the genome, using the Sbfl restriction enzyme. A total of 380 individuals were sequenced: two F_0_ full-sib parents, two F_1_ full-sib offspring, and 376 inbred F_2_ offspring (197 Wt, 178 Cw). We sequenced the 376 samples to ~30x, and as part of a pilot study also sequenced a further plate at higher depth (~100x), which included the F0 and F1 samples. Library preparation and sequencing on the Illumina HiSeq 2000 platform with single-end 100bp reads was performed by Floragenex (Oregon, USA).

Stacks (v 2.6.0) was used to demultiplex libraries, and BWA-mem (v 0.7.17) (55) to align the sequences to the *T. oceanicus* reference genome (Zhang et al. in review) with default parameters. Variant discovery was performed using the *gstacks* utility within Stacks (56), with a minimum mapping quality score of >=20. The resulting catalog was filtered to retain only SNPs with a minor allele frequency > 0.1 and genotyped in > three-quarters of samples in both phenotypes, then converted to vcf format. Genotypes with quality scores less than 20 were filtered using vcftools (v 0.1.16) (35). We rescored X-linked heterozygote calls for each of the male samples as homozygote for the dominant allele if a binomial test revealed significant (P<0.05) differences in allelic depth, and removed heterozygote calls without significant allelic imbalance. We removed genotype calls at loci with sequencing depth < one-third or > three-times the respective sample’s mean sequencing depth, atop a fixed minimum sequencing depth of 10x. Sequencing depth filters were halved for X-linked loci in male samples. Because the presence of PCR duplicates in single-end RAD-seq data is expected to exaggerate heterozygosity, all heterozygotes with strong skew in allelic depth (binomial test P-value < 0.01) were rescored as homozygotes. Association tests between SNPs and Cw presence/absence were performed using Fisher’s exact tests in PLINK (v 1.9) (57), discarding any loci missing in >= 30% of samples, and samples lacking genotypes at >= 60% of loci. P-values were adjusted for multiple testing using Bonferroni correction. A small number (N = 3) of highly significantly Cw-associated SNPs located on Chr4 were removed from the analysis, because visualisation of linkage patterns indicated they showed much stronger linkage with the 0:80 MB region of Chr2 than nearby regions of Chr4, indicating these SNPs were in fact located on Chr2.

#### Whole genome sequencing data

To further investigate candidate Cw-associated regions from the RAD-seq data, we used whole genome sequencing data at ca. 20X depth obtained from wild DNA samples by Zhang et al. (2021). We aligned reads to the genome using bwa-mem2 with default parameters, then removed secondary alignments and PCR duplicates as in Zhang et al. (2021). We used bcftools (58) to call variants on Chr2, which were filtered with vcftools to remove genotypes with sequencing depth < 10 or > 120, mapping quality < 20, and minor allele frequency < 0.05. We used PLINK (v 1.9) to perform principal component analysis after merging the vcf for these samples with the vcf for the RAD-seq samples. To quantify linkage, we first thinned the filtered vcf using vcftools to retain 1% of variants (--thin 0.01). The resulting data were converted to .PED format using PLINK and used as input for Haploview v4.2 (59).

We collected whole sequencing data in 2021/2022 from the same population (Oahu.CC), as well as two other populations (Kauai.CG and Hawaii.UH) in which we observe Cw phenotypes. For these samples, we had recorded whether each individual visibly expressed the curly-wing phenotype. We extracted genomic DNA from and sequenced 90 males (30 from each population) on an Illumina NovaSeq S4, generating 2x150 bp reads at ca. 15X average depth. Library preparation, sequencing, and trimming of adapter and low quality sequences was performed by the Centre for Genomic Research at the University of Liverpool. We aligned reads to the *T. oceanicus* genome V2 with bwa-mem2 and default parameters, and marked PCR duplicates using PicardTools (Broad Institute, 2019). We identified and called variants on Chr2 for each population using bcftools, filtered with vcftools to remove genotypes with sequencing depth < 5 or > 100, mapping quality < 20, and minor allele frequency < 0.1. Linkage was calculated in Haploview after thinning the VCF to retain 0.1% of SNPs. VCFs for each of the population were merged using bcftools, and converted to BED format using Plink. PCA was performed with Plink, excluding SNPs with minor allele frequency < 0.15 and which were not genotyped in >= 80% of samples. Association tests were performed using GEMMA, excluding positions genotyped in less than 90% of samples, and SNPs with minor allele frequency < 0.15. We first created a centred relatedness matrix from variants on Chr2, then ran a linear mixed model with a predictor of Cw phenotype, and the relatedness matrix as a covariate.

Variant calling, linkage, and association tests between Nw and Fw phenotypes were performed as above, except restricting variant calling to the X chromosome, specifying that all samples were haploid (as males only carry a single X), with a maximum sequencing depth of 70, and excluding positions genotyped in less than 85% of samples. For the Oahu.CC population, we combined data collected in the current study (N=30 samples) with that previously collected by Zhang et al. (2021) in 2017, for a total sample of 50. For the Kauai.CG population, we did not include the relatedness matrix as a covariate, given the small sample size and limited statistical power.

#### RNA sampling

We sampled RNA from developing forewing wing buds from a polymorphic F_2_ full-sib family. This F_2_ family resulted from a different F_1_ cross than that used for the generation of RAD-seq data, though it was from the same F_1_ full-sib family. We sampled wing buds at the final instar, immediately prior to adult eclosion, two days after the penultimate moult. We sampled at this stage because curly-wing morphology is not yet externally visible. Thus, at the time of sampling we were unaware of each male’s adult phenotype, so we sampled only one developing (dorsal-right) wing bud and recorded the phenotype of the other wing after adult eclosion.

We retained only unambiguous Cw and Wt phenotypes. Two wing buds from different crickets were pooled per library. For Cw individuals, we retained only males with curliness scores of 2 or 3 for their left wing. For each pooled Cw sample, one male had a curliness score of 2 and the other a curliness score of 3. Flatwing was polymorphic in the inbred F_2_ family used for sampling wings, so we included both male wing vein phenotypes to produce 4 mRNA libraries for each combination of Wt/Cw x Nw/Fw. RNA sampling was performed under CO_2_ anaesthesia to minimise effects of surgical wing bud removal on welfare and gene expression, and developing wing tissue was removed using dissection scissors. Wing tissues were submerged in RNAlater and stored at -20° C.

#### RNA-seq processing and differential expression analysis

mRNA libraries were prepared using PolyA selection and sequenced on an Illumina NovaSeq S1. Reads provided after trimming adapter and low quality sequences by the Centre for Genomic Research, University of Liverpool. We filtered sequences with similarity to known Eukaryotic ribosomal RNAs using sortmerna (60). Filtered reads were aligned to the genome using HISAT2 (61), then individual transcriptomes assembled and merged using Stringtie (62). Transcript abundances were subsequently quantified at the ‘gene’ level. We filtered any genes not expressed at greater than one count-per-million in at least four samples. After filtering, 12,053 genes were retained for analysis.

Differential expression analysis was performed in DEseq2 (63), with models tested using likelihood ratio tests and an FDR-adjusted P threshold of 0.05 for a gene to be considered significantly differentially expressed. Visualisation of gene expression patterns revealed that two samples, one CwFw and one CwNw male, showed gene expression patterns discordant with the other samples, and were therefore removed from the gene count matrix for the differential expression analysis (though we confirmed that these samples showed consistent patterns of expression in differential expression comparisons). To assess functional enrichment, we first used blastx with an e-value filter of 1e^-6^ to identify orthologs in *Drosophila* for all genes in the filtered transcriptome. Next, we performed binomial tests between the number of genes in each category in differentially expressed (using a relaxed threshold of P_adj_<0.1 to increase statistical power) and reference (all genes in the filtered transcriptome) gene sets using PANTHER (64), with Bonferroni-adjustment of P-values to account for multiple testing.

#### Mate preference trials

Males and females (83 of each sex) were isolated from the Oahu.CC stock population at the final instar prior to adult eclosion to ensure virginity. Cw and Fw phenotypes are both present and polymorphic in this population. Individuals were phenotyped at adulthood, and used in mate preference trials at 5-10 days post-adult eclosion, when both sexes are reproductively mature. To conduct trials, a male and female were chosen at random and placed in a 210 × 230 mm arena together under red light and then left to interact for 10 minutes. Trials were filmed on a Nikon D3300 digital camera. We recorded whether the male produced (or attempted to produce) courtship song, and whether the female mounted the male within the 10 minute trial.

Trial data were analysed using GLMs with binomial error distribution in R v4.0.2 (R Core Team 2020): female mount (Y or N) ~ male court (Y or N) ⨉ male morph (WtNw, Fw [CwFw & WtFw pooled due to small sample size], or CwNw). Binomial GLMs were checked for overdispersion and tested using type III Wald’s Chi-squared tests. Although we originally used each male and female in multiple trials (up to four times), having prevented them from spermatophore transfer during trials to ensure virginity, attempts to include male and female ID in the model as random effect terms using *lme4* (66) produced convergence errors, so we used only the first trial for each male and female to avoid pseudoreplication and confounding effects of prior experience in our analyses.

#### Life history assays

To test for correlated fitness effects of Cw in non-wing tissues, we recorded structural size (pronotum length to nearest .1 mm) and wet mass (to the nearest .01 mg) at adulthood for 91 males and 48 females from a mixed stock population. Mass and survival status were recorded every 7 days up to 91 days post-adulthood in females and 84 days post-adulthood in males. While mass can vary with age, pronotum length is fixed at adult eclosion. Therefore, we measured pronotum length 3 times and used the mean average to account for measurement error. All crickets were virgin, which we ensured by rearing them in isolation. We used pronotum length and mass measures to calculate scaled mass index (SMI) separately for each sex, a measure used to estimate body condition (43). Survival was recorded up to a maximum of 91 days in females, and 84 days in males.

We used mass measurements to examine adult size and SMI to examine body condition; the latter incorporating information about both mass and structural body size (pronotum length). Analysis of female mass across adult lifespan was performed using a linear mixed model using *lme4* in R, including linear and quadratic terms for adult age in days, Wt/Cw phenotype, and interactions. For males, there was no evidence of a relationship between age and mass, so we analysed mass at 14-days post-eclosion, an age at which males are reproductively active but are less likely to show signs of senescence. We used a linear model with predictor terms of wing shape (Wt/Cw) and wing venation (Nw/Fw), and removed the interaction between the two because it did not approach statistical significance. Note that Fw/Nw could not be phenotyped in females as Fw expression is male-limited. For models with and without interactions we used Type II and III Wald’s Chi square tests, respectively, using the *car* package in R.

Lifetime survival probability was analysed using Cox proportional hazards regression in the R package *survival* (67). We included predictors of wing shape and SMI for females, and wing shape, wing venation and SMI for males. Interactions that did not approach significance were removed from the model.

#### Cricket rearing and husbandry

Unless otherwise indicated, analyses relate to samples derived, or collected, from a wild population at a Community Center in Manoa, Oahu (Lat.: 21.316219, Long.: -157.809922, Fig. 1B). This is the population in which we first observed the Cw phenotype in 2017. Since it was first observed, Cw has been stably present at approximately 50% frequency in the Community Center population (personal observations, JGR & NWB). We also include whole genome sequencing samples from two more populations in which Cw and/or Fw phenotypes also co-occur: the Kilauea Common Ground on the island of Kauai (Lat.: 22.197576, Long.: -159.417606), the University of Hawaii Hilo campus on the island of Hawaii (Lat.: 19.703186, Long.: -155.080237).

All experiments were performed using a stock population, or inbred populations derived from the stock population, derived from eggs of wild caught *T. oceanicus* females captured in 2017 from a location in Manoa, Oahu, HI. Eggs from wild-caught females were transported back to the University of St Andrews, UK, and were reared as a mixed stock population across multiple 20L boxes at 25c, on a 12:12 photoreversed light:dark cycle. Crickets were fed Purina Rabbit Chow (Burgess Excel) and provided with cotton pads moistened with dH2O for water and oviposition, as well as cardboard shelter.

#### Extraction of nucleic acid samples

Purified DNA and RNA samples were extracted as previously described in Zhang et al. (2021). Briefly, a CTAB/chloroform-based extraction protocol was used to extract DNA from neural or leg tissues. RNA samples were extracted using a Trizol/chloroform-based protocol, with subsequent washes performed following the ThermoFisher Purelink protocol.

### Supporting figures and tables

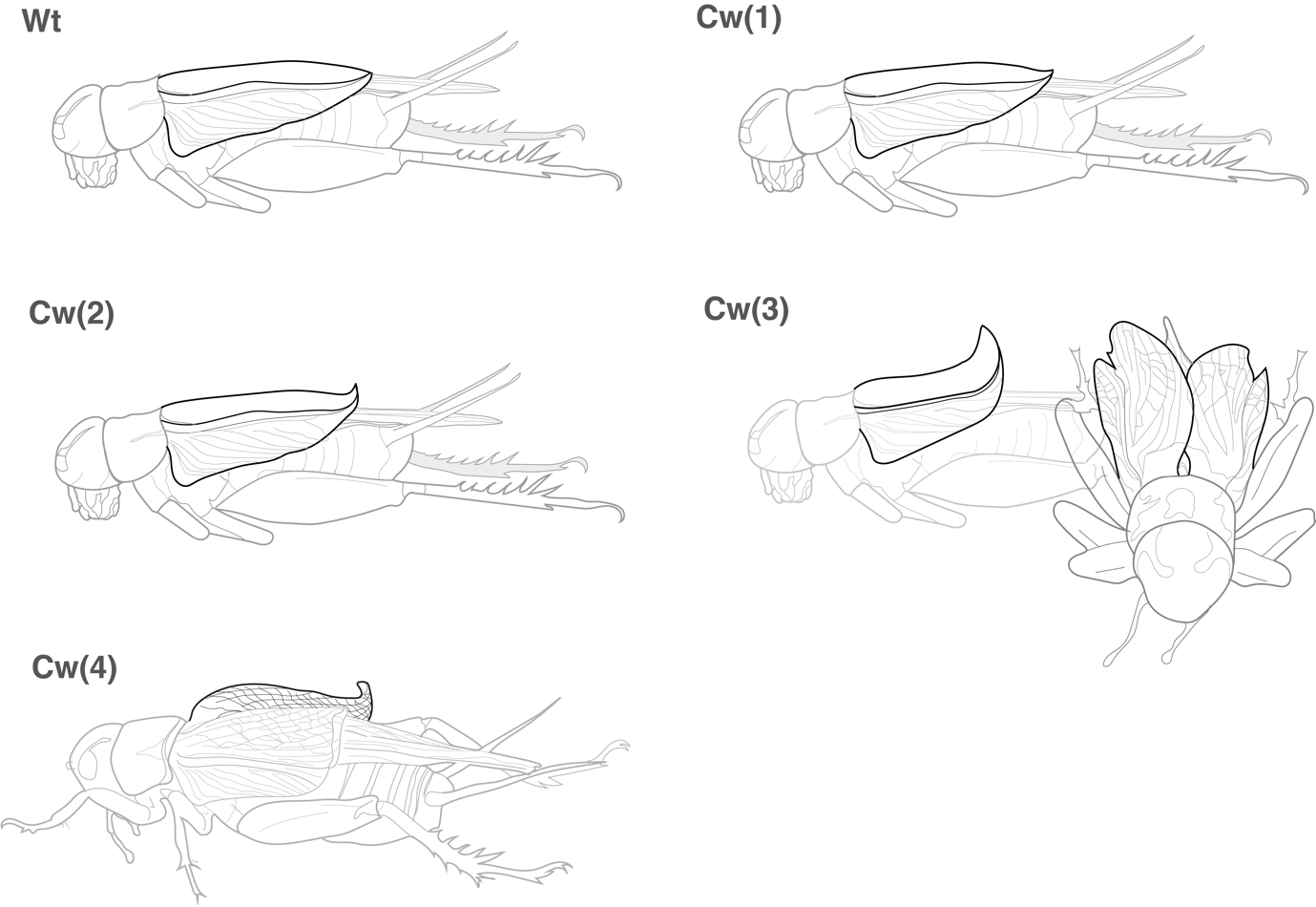

**Figure S1**. Illustrations of Wt and Cw morphology, with examples of wings with curliness scores of 1 (slight) to 4 (extreme).

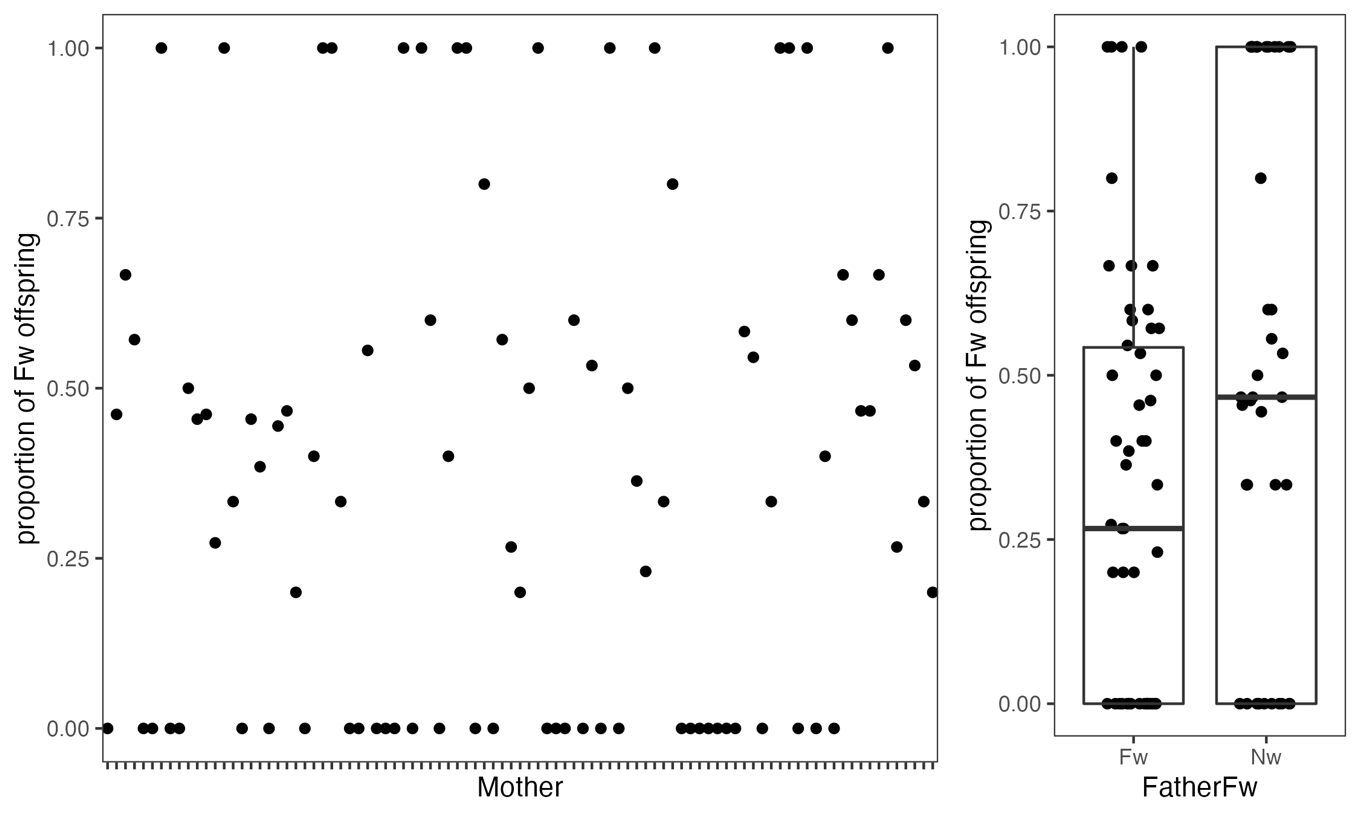

**Figure S2.** Left: proportions of Fw male offspring across families. Fw tended to be present at 0%, 100% or ca. 50% frequencies, suggesting heritable variation and consistent with expectations of a simple Mendellian trait. Right: proportions of Fw males produced by crosses involving Fw and Nw fathers, illustrating the lack of an increased proportion of Fw males sired by Fw fathers, supporting X-linked inheritance.

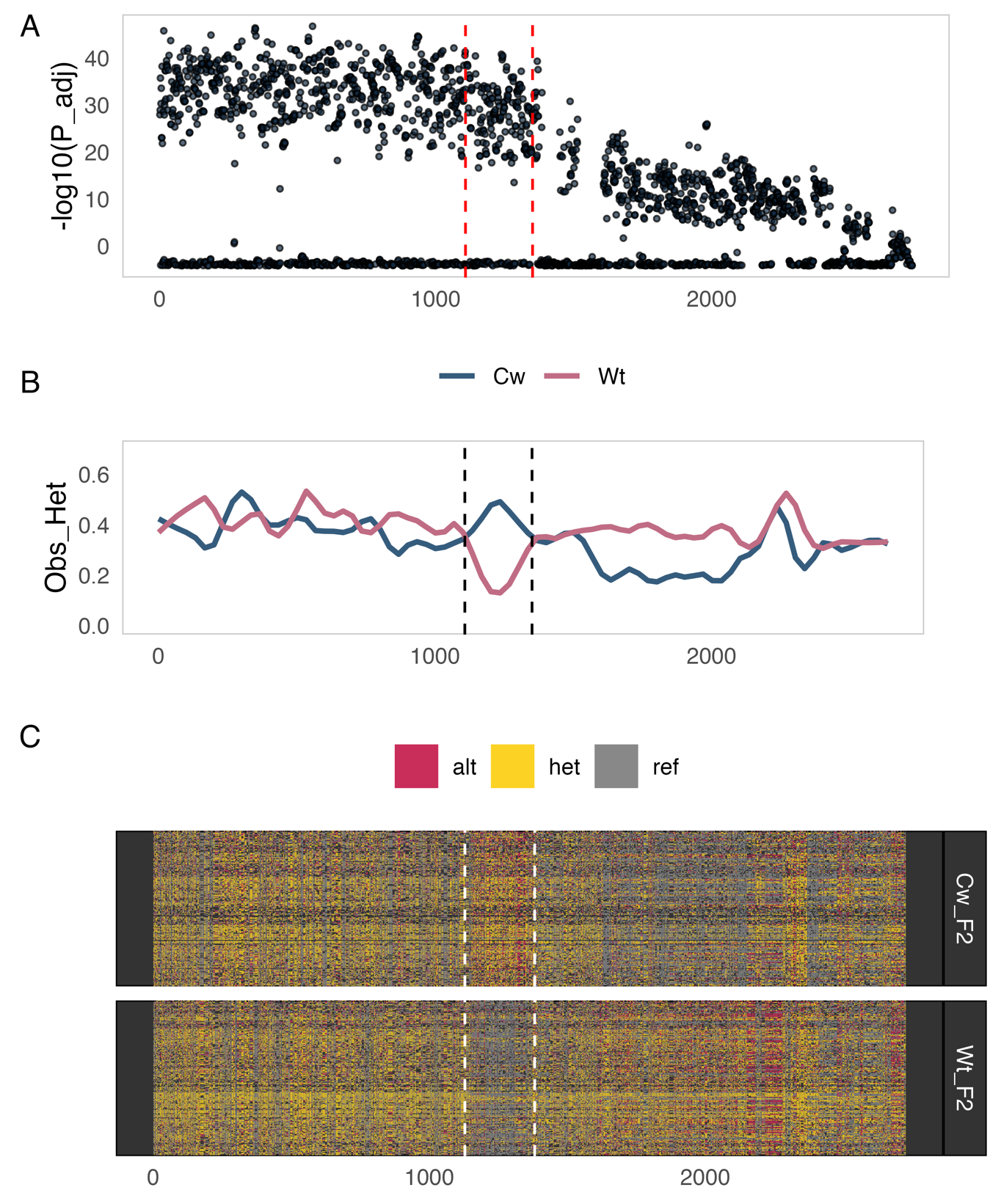

**Figure S3.** Patterns of **A)** genetic association with Cw; **B)** heterozygosity; and **C)** genotype across SNPs (not scaled by genomic coordinates) across SNPs located on Chr2. Dashed lines indicate the region between 66 and 83 MB which seemed to be highlighted in panels B and C.

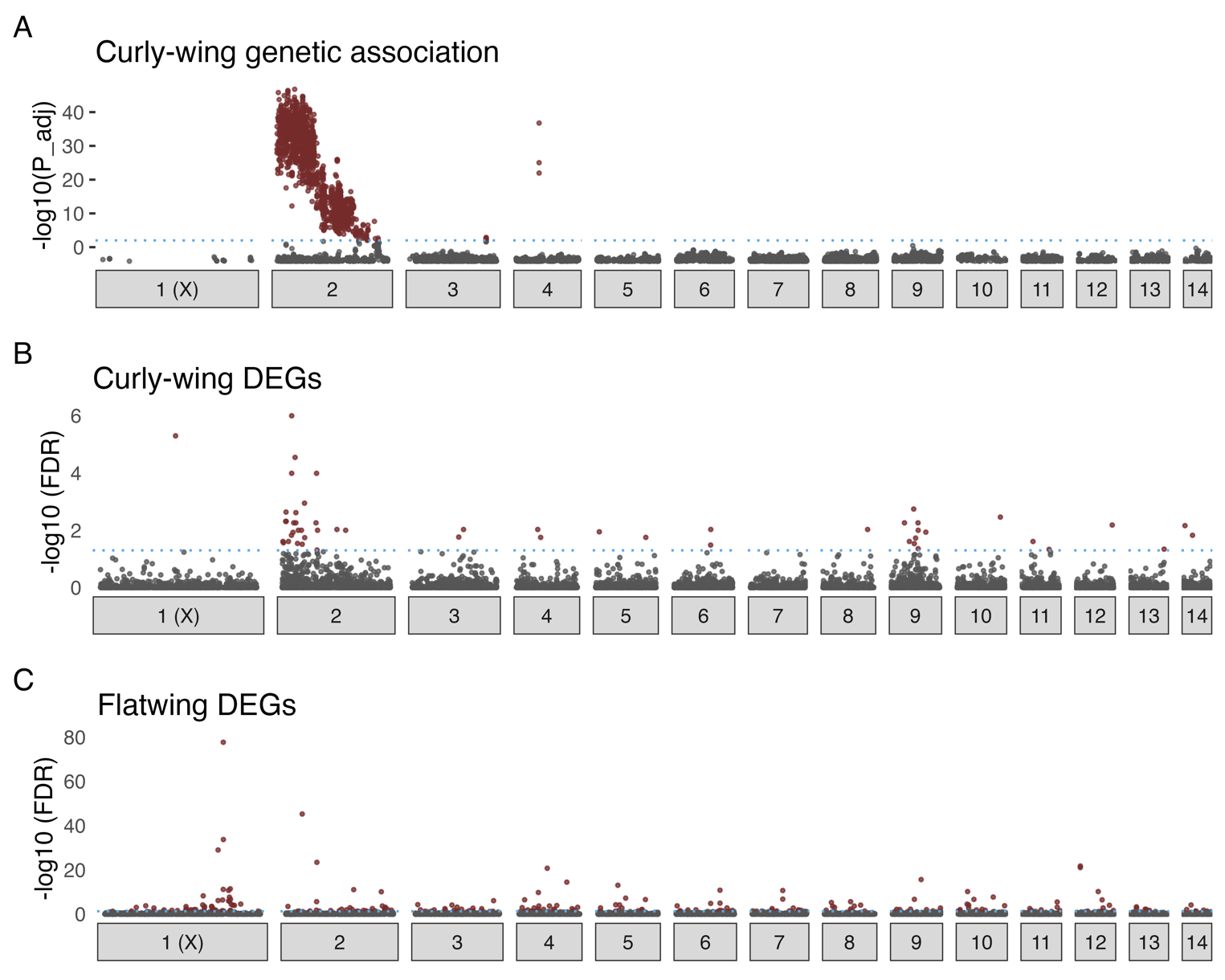

**Figure S4.** (**A**) Genome-wide association significance (-log_10_ Bonferroni-adjusted P) between allelic variants and Cw phenotype. (**B,C)** Locations and relative significance of genes DE between (B) Wt and Cw phenotypes, for which the causative region is on chromosome 2, and (C) Nw and Fw phenotypes, for which the causative region is located on the X chromosome. Note the order of magnitude difference in Y-axis scales between B and C. Pink labels indicate the chromosome harbouring the causative genetic variant(s) for each wing phenotype, based on genomic association. In all plots, points are coloured by statistical significance (A: P_adj_<0.01; B,C: FDR<0.05) also indicated by the dotted horizontal line. Illustrations visualise curly-wing and flatwing phenotypes in the respective panels.

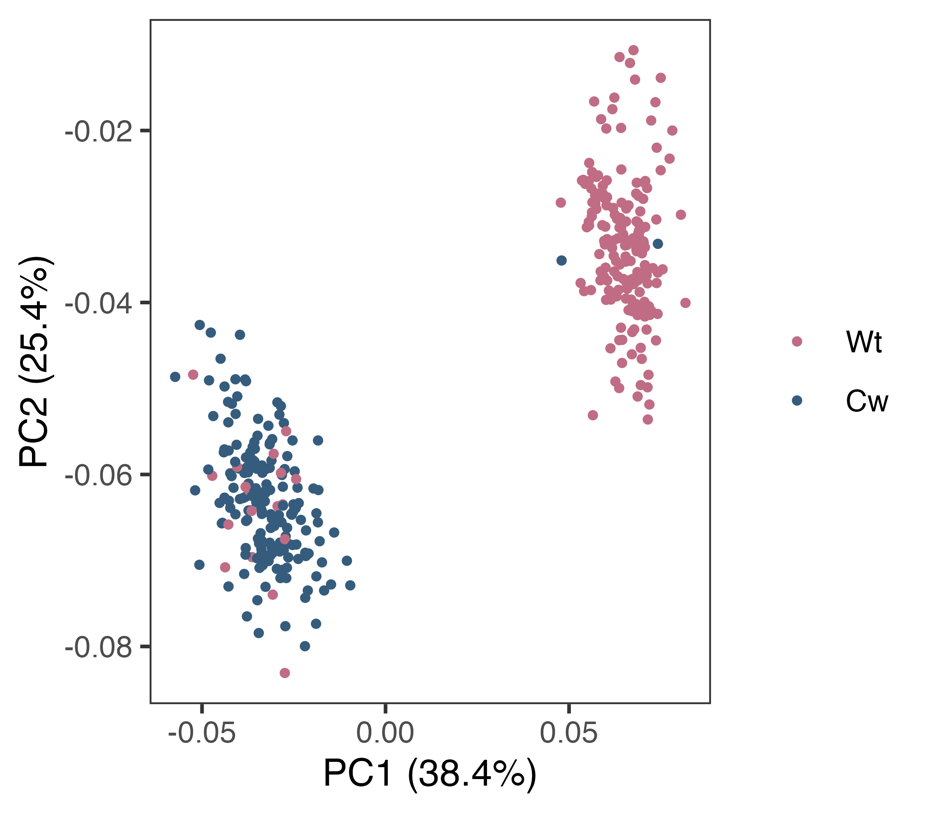

**Figure S5.** PCA generated using RAD-seq SNPs from the region of 0 to 80 MB on chromosome 2.

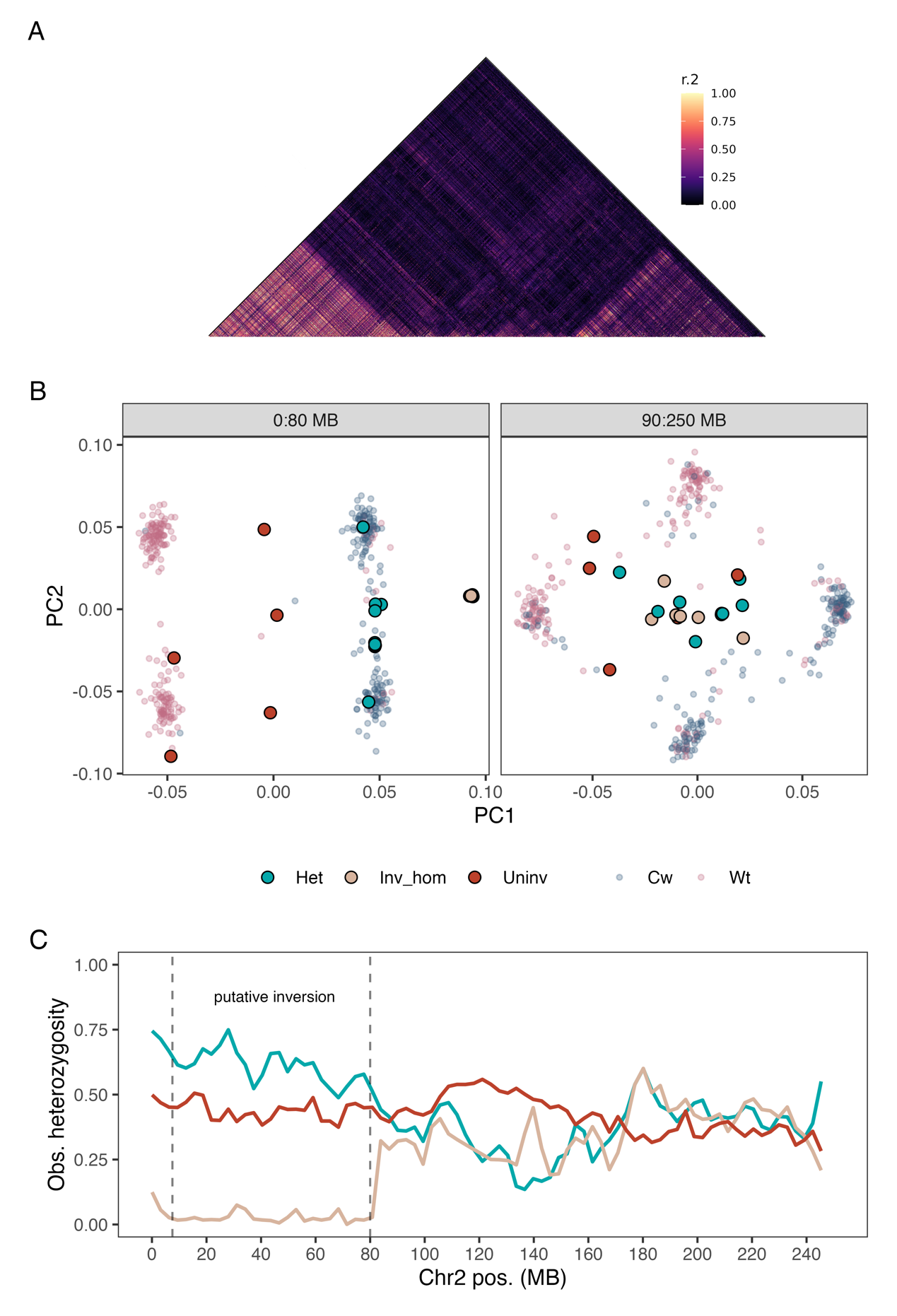

**Figure S6. Evidence of a large inversion in the region of Chr2 associated with curly-wing. (A)** Strong linkage between SNPs in the region of the candidate inversion on Chr2. **(B)** PCA incorporating RAD and WGS data for variants present in both filtered datasets on Chr2, including SNPS within the Cw-associated region identified by the RAD-seq analysis (left), and across the remainder of the chromosome (right). Small circles show RAD-seq samples coloured by phenotype, whereas larger filled points show WGS samples, filled by putative genotype with respect to the candidate inversion. **(C)** Heterozygosity on Chr2 between samples from the three putative genotypes, with the candidate inversion highlighted with vertical dashed lines bracketing the first 80 Mb. Solid lines show LOESS trended means.

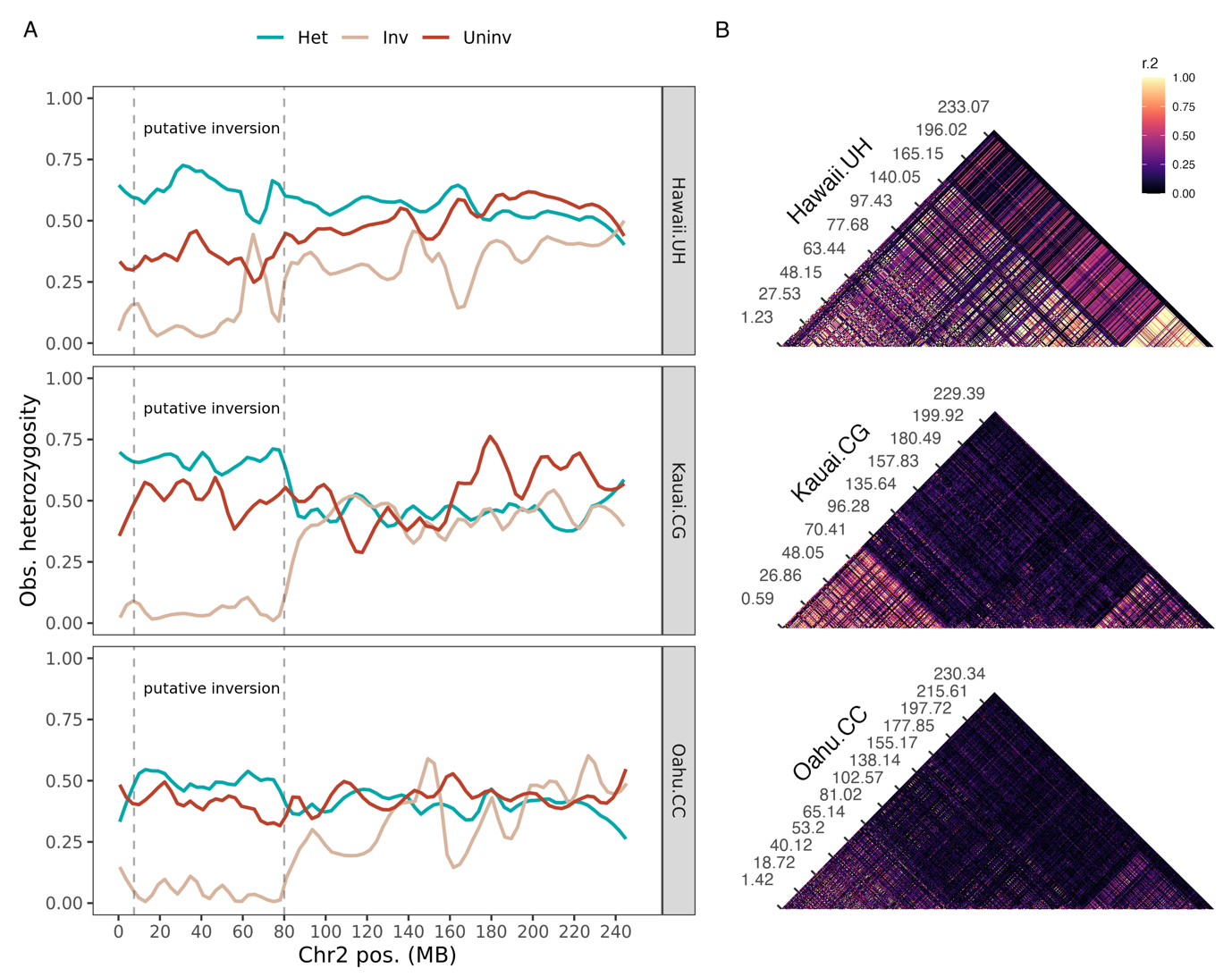

**Figure S7. A)** Heterozygosity between inferred genotypes (based on PC1 values from Fig 4A) across the putative inverted region of 7.5:80 Mb on Chr2, in three populations. **B**) Linkage between thinned (0.0005) SNPs genotyped in >50% of samples across all three populations.

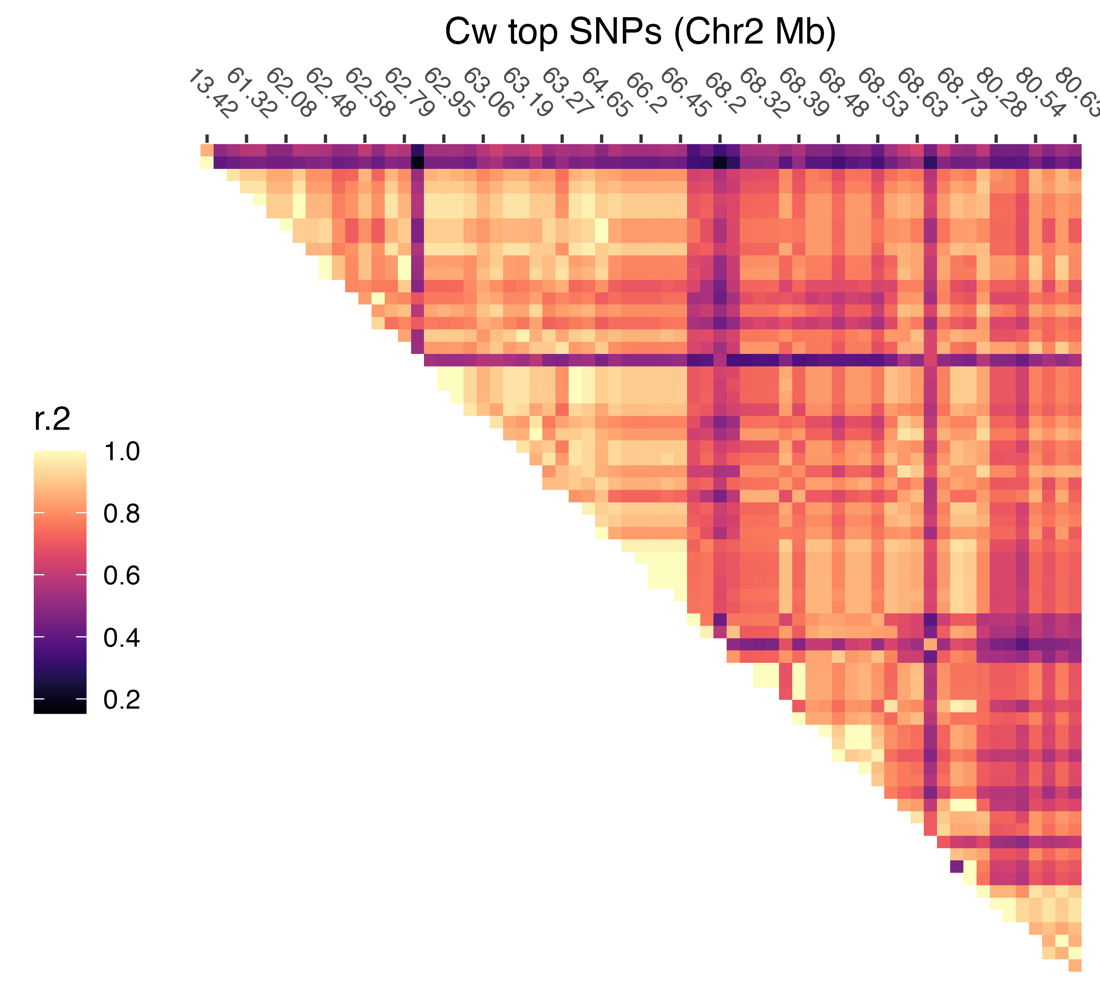

**Figure S8**. Linkage between the top 100 Cw-associated SNPs.

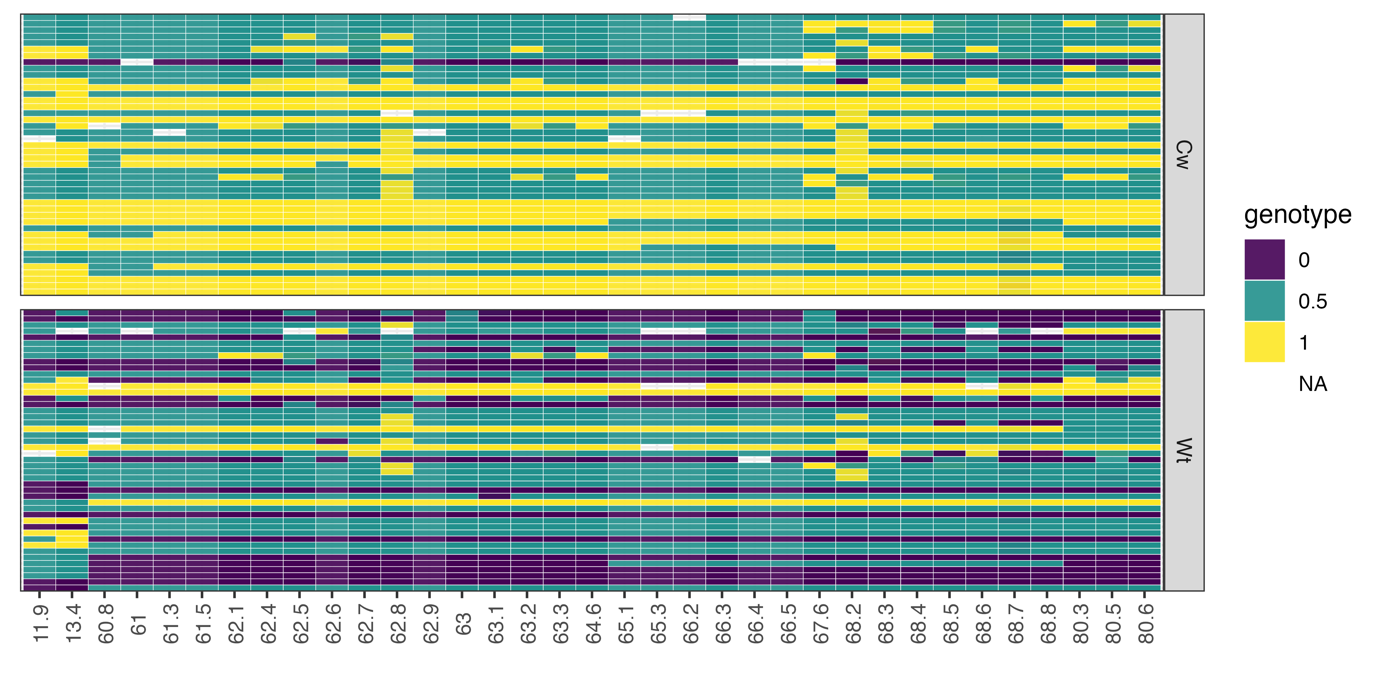

**Figure S9**. Genotypes (0=homozygous major allele; 0.5=heterozygote; 1=homozygous minor allele) across samples for each of the top 100 Cw-associated SNPs.

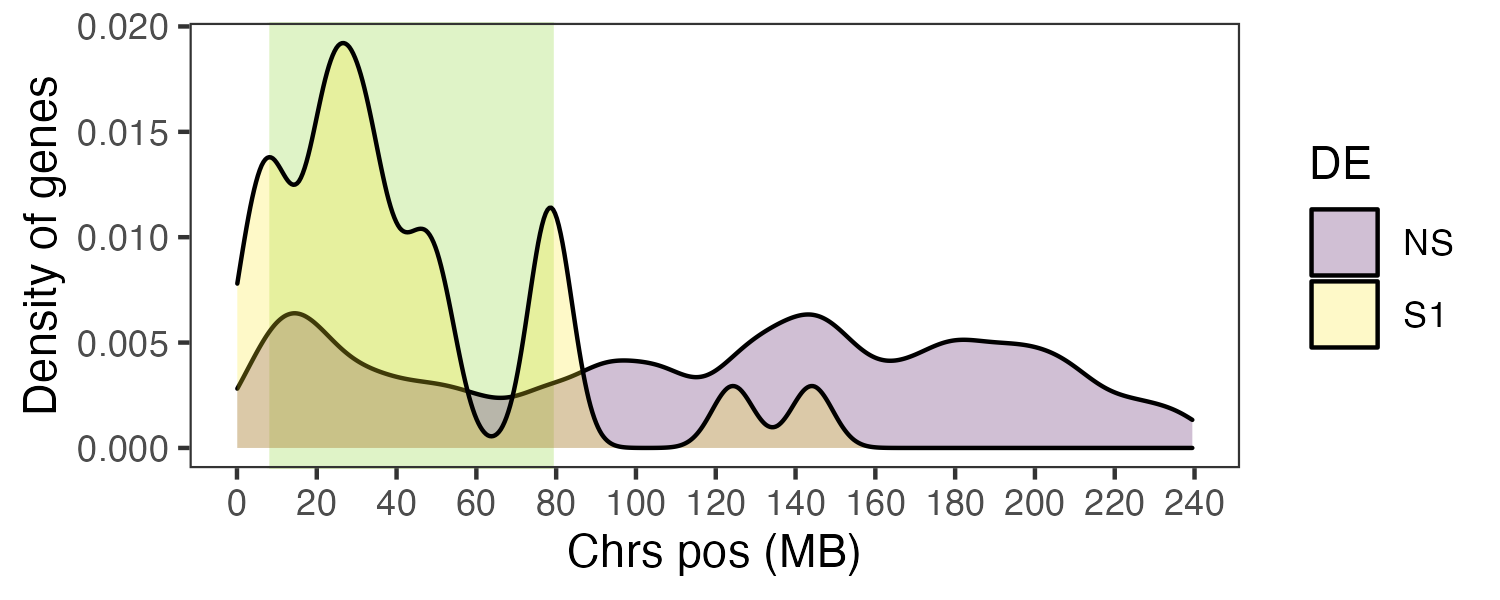

**Figure S10.** Density of expressed genes along Chr2, with DE_Cw_ genes shown in yellow.

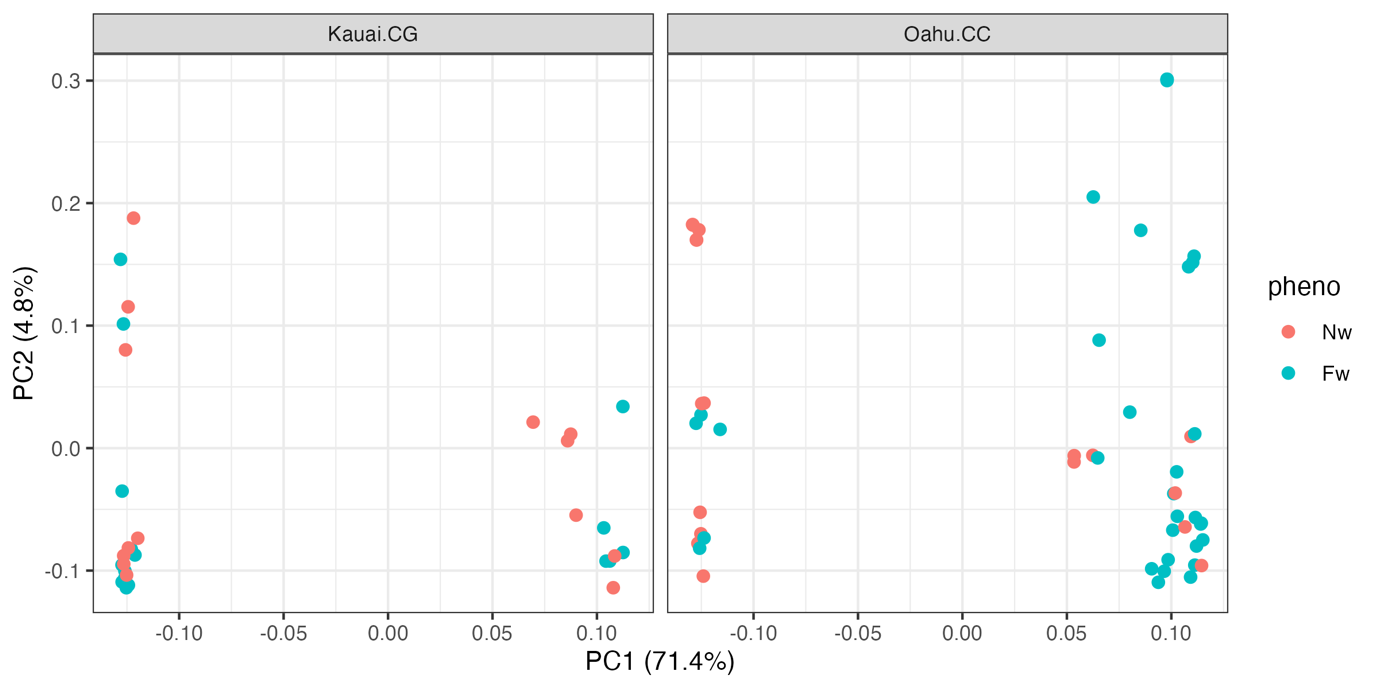

**Figure S11.** PCA of variants on Chr1 (the X) using samples from Oahu.CC (N=50) and Kauai.CG (N=30). Samples were clearly separated into two clusters, but cluster was not obviously associated with Fw/Nw phenotypes.

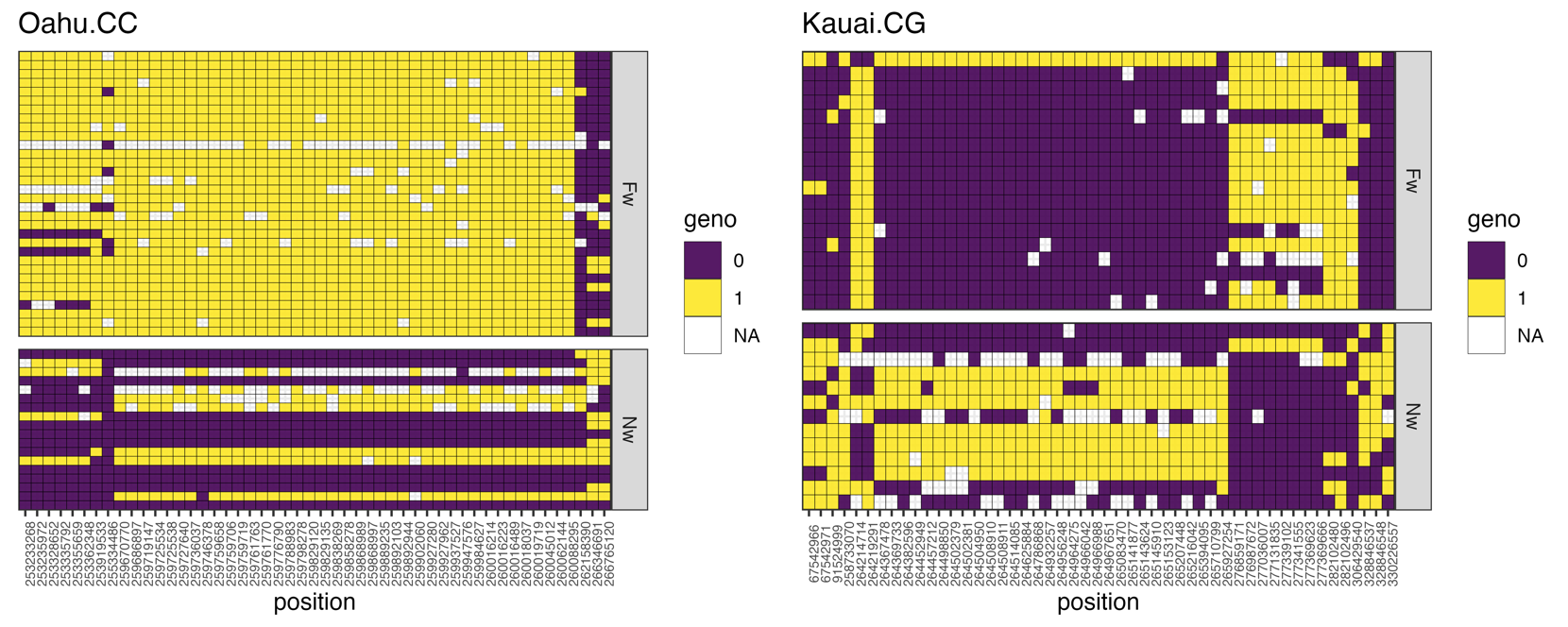

**Figure S12.** Genotypes across samples for each of the top 50 Fw-associated SNPs (outside of the region of 95.7 – 253.2 Mb on the X) for Oahu.CC and Kauai.CG populations.

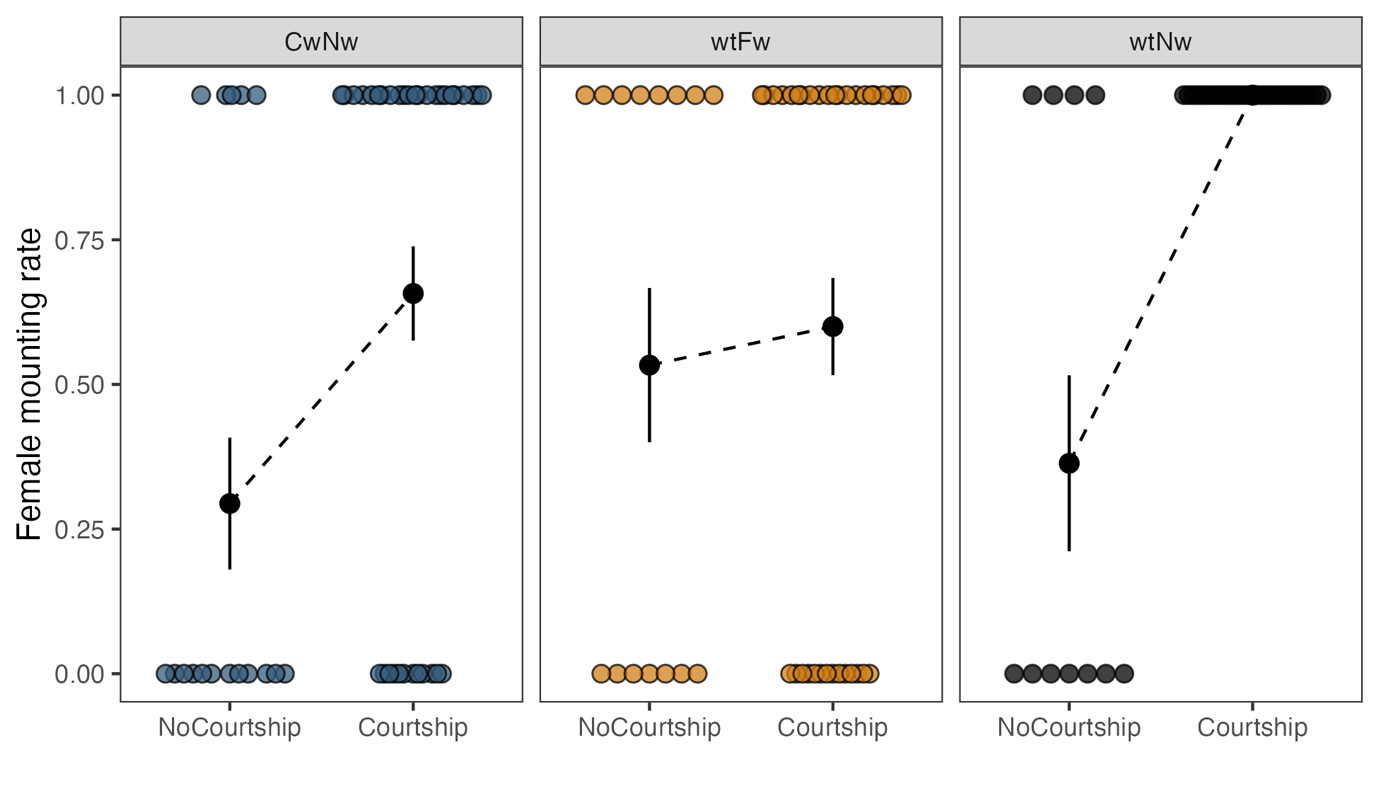

**Figure S12.** Effects of courtship (measured by attempt to produce courtship song) by males of different wing phenotypes upon rates of female mounting, including repeated measures (cf. Fig. 7).

**Table S1.** Criteria used for scoring wing curliness on a quantitative scale.

| Score | Description |
| --- | --- |
| 0 | None; no visible curliness |
| 1 | Slight; curliness visible only on close inspection (wing tips or edges slightly upturned) |
| 2 | Moderate; readily visible but not striking. Greater than 3/4 of the wing surface retains an ordinary shape. |
| 3 | Striking; wing is furled (curls past 180 degrees), or makes little contact with the body (between 1/4 and two-thirds of the wing makes no contact with the body). |
| 4 | Extreme; the wing makes almost no contact with the body of the cricket (more than half of the wing makes no contact with the body). |

**Table S2.** Results of a binomial GLMM testing for an association between expression of Cw and expression of Fw among male offspring (N=1,220), including rearing density as a covariate, and a nested random effect term representing heritability (1|cross_group/Father/Mother).

| Response |  | X_1_^2^ | P |
| --- | --- | --- | --- |
| Cw expression | Fw expression | 0.545 | 0.460 |
|  | Rearing density | 6.291 | 0.012 |

**Table S3**. Results of Cox proportional hazards regression of female and male longevity associated with wing phenotype and scaled mass index (SMI).

| Response | Predictor | N | Hazard ratio | 95% CI | P |
| --- | --- | --- | --- | --- | --- |
| Female longevity | wing_shape | 46 |  |  |  |
|  | Cw |  | — | — |  |
|  | Wt |  | 0.49 | 0.39, 0.60 | <0.001 |
|  | SMI | 46 | 0.99 | 0.98, 1.0 | <0.001 |
| Male longevity | wing_shape | 88 |  |  |  |
|  | Cw |  | — | — |  |
|  | Wt |  | 0.86 | 0.39, 1.93 | 0.7 |
|  | wing_veins | 88 |  |  |  |
|  | Fw |  | — | — |  |
|  | Nw |  | 0.61 | 0.23, 1.59 | 0.3 |
|  | SMI | 88 | 1.01 | 0.99, 1.02 | 0.4 |

**Table S4**. Results of a linear mixed model for female mass (across adulthood, with random intercepts for stock box and ID nested within stock box), and linear model for male mass (at 14 days post-adulthood).

| Response | Predictor | X_1_^2^ | P |
| --- | --- | --- | --- |
| Female mass | days | 200.232 | <0.001 |
|  | days^2^ | 81.045 | <0.001 |
|  | wing_shape | 5.009 | 0.025 |
|  | days:wing_shape | 8.409 | 0.004 |
|  | days^2^:wing_shape | 11.380 | <0.001 |
| Male mass | wing_shape | 1.225 | 0.268 |
|  | wing_veins | 1.247 | 0.264 |

**Table S5.** Results of linear models for female SMI (across adulthood, with random intercepts for stock box and ID nested within stock box), and males (at 14 days post-adulthood).

| Response |  | X_1_^2^ | P |
| --- | --- | --- | --- |
| Female mass | Intercept | 878.224 | <0.001 |
|  | days | 164.907 | <0.001 |
|  | days^2^ | 54.955 | <0.001 |
|  | wing | 0.418 | 0.518 |
|  | days:wing | 2.716 | 0.099 |
|  | days^2^:wing | 3.497 | 0.061 |
| Male mass | Intercept | 1346.646 | <0.001 |
|  | wing_shape | 1.219 | 0.270 |
|  | wing_veins | 0.324 | 0.569 |

**Table S6**. Description of typical singing-capable and adaptive reduced-song *Teleogryllus oceanicus* wing phenotypes.

|  | | **Wing venation** | |
| --- | --- | --- | --- |
|  |  | **Nw** | **Fw** |
| **Wing shape** | **Wt** | WtNw – the ‘typical’ ancestral male phenotype capable of producing song at ordinary levels due to sexually dimorphic, specialised song-producing structures on the forewing, and the proper engagement of the scraper/file mechanism during stridulation | WtFw – male unable to produce song at normal amplitude due to the reduction of sound-producing structures on the forewing, including a strongly reduced stridulatory file |
|  | **Cw** | CwNw – male unable to produce song at normal amplitude due to unusually curled wings precluding proper engagement of the scraper/file mechanism during stridulation | CwFw – male unable to produce song at normal amplitude due to a combination of the reduction of sound-producing structures on the forewing, and unusually curled wings that preclude proper engagement of the scraper/file mechanism during stridulation |
